# Activation pathway of the human voltage-gated proton channel

**DOI:** 10.64898/2026.10.06.757015

**Authors:** Mingzhe Shen, Daniel Kortzak, Christophe Jardin, Shuo Han, Nathaniel Felbinger, Shizhen Wang, Brian G. Pierce, Boris Musset, Thomas E. DeCoursey, Jana Shen

**Author notes:** Joint first author (Mingzhe Shen, Daniel Kortzak).

## Abstract

Many protein functions are coupled to proton transfer, yet the underlying mechanisms remain extremely difficult to resolve by experiment or conventional molecular dynamics (MD) simulations. The human voltage-gated proton channel H_V_1 (hH_V_1) plays key roles in human physiology, but how voltage and pH together drive its activation remains unresolved. Experimental structures of hH_V_1 are limited to the X-ray structure of a mouse chimera and NMR models of a truncated hH_V_1, both representing an intermediate resting state. Here we determine the activation pathway of hH_V_1, together with structural models of its resting and activated states, using an approach that integrates all-atom continuous constant-pH MD (CpHMD), weighted-ensemble (WE) path sampling, and AlphaFold modeling. Starting from an AlphaFold model selected for consistency with resting-state experimental data, over 100 *µ*s of simulations reveal a mechanism in which transmembrane voltage, assisted by proton titration, drives an outward displacement of the transmembrane segment S4 accompanied by a one-click reorganization of countercharge pairings. These events are enabled by a delicately balanced electrostatic landscape and pH-modulated breathing motions in the resting state that prime the channel for protonation-state switches upon voltage sensing, a principle that may underlie other proton-coupled voltage-gating processes. The simulations also uncover lipid-binding sites at which POPG selectively stabilizes the S4-up conformation, whereas POPE can stabilize either state. These predictions are validated against extensive electrophysiology, mutagenesis data, and new smFRET experiments, which support the one-click S4 displacement and the lipid effects. WE-CpHMD, implemented in the AMBER package, is applicable to other proton-coupled processes that remain elusive.

## INTRODUCTION

Voltage-gated proton channels H_V_1^1–3^ are proton-selective ion channels that extrude acid from cells in response to membrane depolarization or intracellular acidification.^4^ H_V_1 has been identified in a variety of cell types across different species.^5^ In humans, H_V_1 (hH_V_1) plays physiological roles such as immunity, reproduction, and cancer.^5,6^ In stimulated phagocytes, reactive oxygen species produced by NADPH oxidase depolarize the membrane and acidify the cytoplasm; these effects are countered through activation of hH_V_1.^7^ During sperm activation, hH_V_1 extrudes protons to raise intracellular pH, triggering the vigorous motility required for egg fertilization.^8^

Unlike other voltage-gated ion channels, in which voltage sensing and ion conduction occur in distinct domains, hH_V_1 conducts protons directly through its voltage-sensing domain (VSD), comprising transmembrane segments S1–S4.^2,3,9,10^ S4 contains the three gating arginines, R1, R2, and R3 (Figure 1a). H_V_1 exists as a dimer in humans and mice, although each monomer is independently functional.^9–11^ Over the past two decades, tremendous effort and progress have been made in experimental (see a recent review^5^) and computational studies of hH_V_1.^12–22^ However, experimental information remains limited. A medium-resolution X-ray structure (PDB: 3WKV)^23^ of a chimeric construct (mH_V_1cc) that combines the mouse H_V_1 sequence, the S2–S3 region of *Ciona intestinalis* phosphatase VSD (Ci-VSD), and the coiled-coil domain of yeast GCN4 offered the first glimpse at a S4-down resting state. This structure, however, is not well suited for structure-function investigation of hH_V_1 due to the difference in the hydrophobic plug or gasket (HG), which forms a dielectric barrier to water and ion permeation.^24^ In hH_V_1, the HG is formed by V109 (S1), F150 (S2), and V177/V178 (S3),^13,25–28^ whereas in mH_V_1cc, I173/V174 (V177/V178 in hH_V_1) are located roughly one helical turn lower than V105 and F146 (V109 and F150 in hH_V_1), with I173 facing lipids. Indeed, the relative positions of F146 and R2/R3 in mH_V_1cc were suggested to differ from those in NMR models of a truncated hH_V_1 construct in an intermediate resting state,^29^ although the side chains of R2, R3, and the selectivity-filter D112 were not resolved.

**Figure 1:**
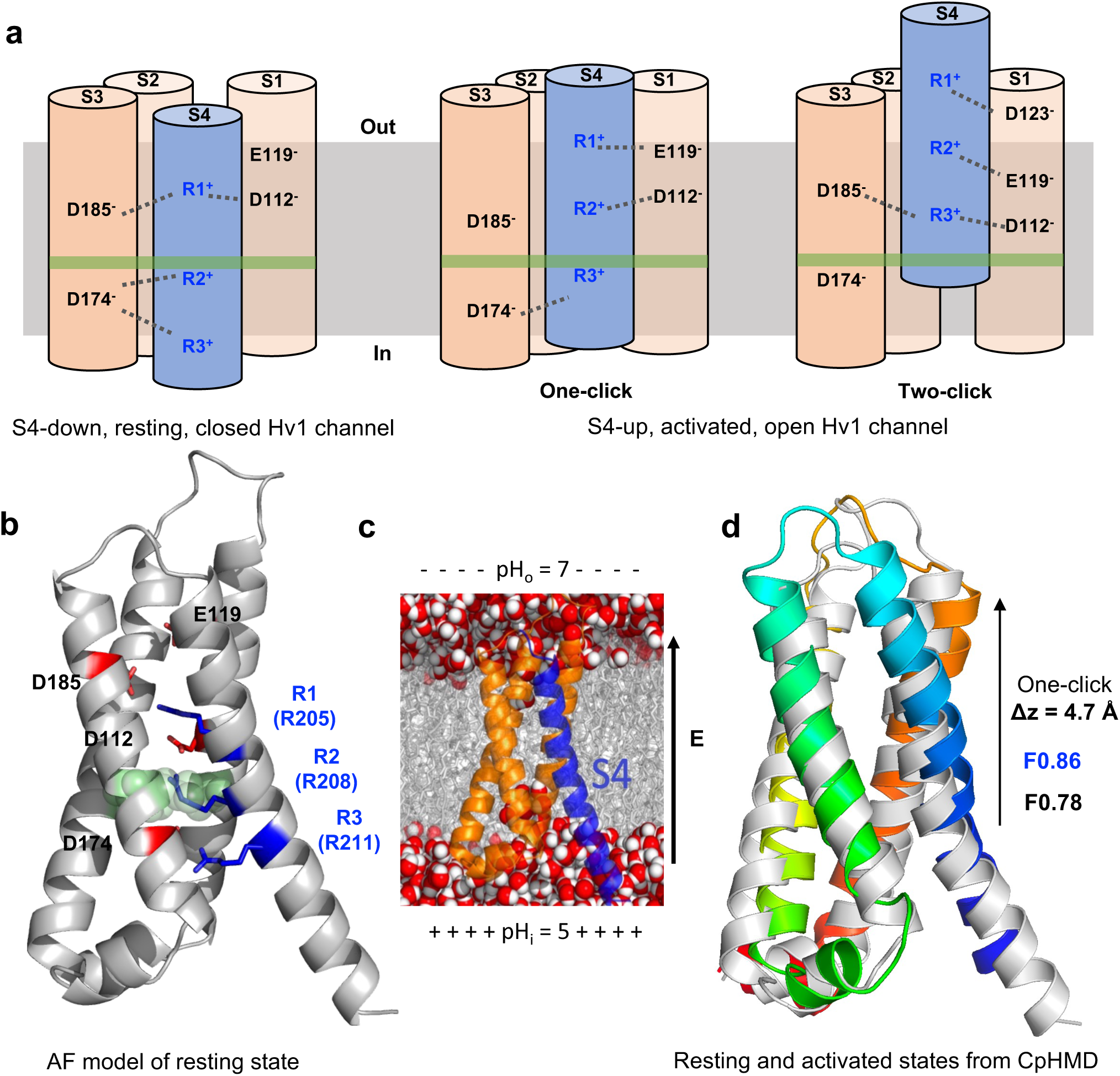
Current models of hH_V_1 activation and predictions by WE-CpHMD simulations. **a.** Proposed resting-state (left) and two alternative activated-state models (middle and right) of hH_V_1. Salt bridges involving the gating arginines and their counter charges are indicated. The green region marks the HG level. **b.** AF model 1 of S4-down hH_V_1 (AF model 2 and Robetta model in Supplementary Figure S1). HG residues are shown as spheres; gating arginines and counter-charges are labeled. **c.** Simulation setup. In the WE-CpHMD simulations, a membrane potential of 200 mV (as a uniform electric field of 0.0583 kcal mol*^−^*^1^ Å *^−^*^1^ e*^−^*^1^) was imposed, while pH_o_ = 7 and pH_i_ = 5 were assigned to residues above and below the HG, respectively. The replica-exchange CpHMD simulations were were performed under symmetric pH and no applied membrane potential. **d.** Overlay of the representative activated (rainbow) and resting (gray) conformations of hH_V_1 aligned on S1 to S3 shows a one-click displacement of S4 by one helical turn (Δ*z* = 4.7 Å), with the S3–S4 angle widening by 2.7*^◦^*. Conformations are cluster centers from the extended WT CpHMD simulations (last 500 ns) for the resting and activated states. Calculated FRET efficiencies for the activated (blue) and resting (black) conformations are shown.

Solvent accessibility data indicate that, upon hH_V_1 activation, R2 but not R3, passes above the HG, gaining access to the extracellular solution.^13,25–28^ Electrophysiology data^13,30–32^ further suggest that the selectivity-filter residue D112 forms a salt bridge with R1 in the resting state and switches to a salt bridge with R2 in the activated state (Figure 1a). Together, these data suggest a R2D, or one-click, model,^13^ consistent with the X-ray structures of resting and activated Ci-VSD^33^ and the EPR data on hH_V_1.^28^

An alternative R3D, or two-click, model – in which R3 moves above the HG and forms a salt bridge with D112 in hH_V_1 (Figure 1a) – was recently suggested by a molecular dynamics study^18^ employing fixed protonation states, hereafter referred to as conventional MD (cMD). This work has several important limitations. First, the simulations were initiated from a resting-state hH_V_1 model based on the mH_V_1cc X-ray structure,^23^ which differs from hH_V_1 in the HG residues (see discussion earlier). As a result, the HG in the derived hyperpolarized hH_V_1 model is disrupted.^18^ Second, protonation states were fixed at the program defaults (deprotonated Asp/Glu and neutral His), and no ΔpH was considered. These structural and methodological limitations cast doubt on the validity of the proposed mechanism.

Beyond the one-vs. two-click debate, important questions regarding the voltage- and ΔpH-gating mechanism of hH_V_1 remain open^34^ (Figure 1a, 1b). Which parts of the channel (e.g., S4 alone, or S4 together with S1) move toward the extracellular side (also referred to as outward or upward) upon channel opening, and what is the magnitude of S4 displacement? What are the salt bridges (or charge-countercharge pairs) stabilizing the resting and activated states? How does S4 motion rearrange these salt bridges to enable proton conduction? How does protonation dynamics facilitate channel activation?

Here we present a computational framework to unravel the activation pathway of hH_V_1. Starting from a resting-state model selected from those generated by the AlphaFold (AF)-based^35^ AF-cluster^36^ approach, GPU-accelerated all-atom particle-mesh Ewald continuous constant pH MD (CpHMD)^37–39^ simulations were conducted under experimentally relevant membrane potential (ΔV_m_) and ΔpH, with the weighted-ensemble (WE) protocol^40,41^ for enhanced sampling of channel-opening events. Over 100 *µ*s MD data depict voltage-gated, proton-coupled conformational activation of hH_V_1, providing a mechanistic framework linking voltage gating, salt-bridge rearrangements, and proton conduction. The WE-CpHMD framework is broadly applicable to proton-coupled channels and transporters that remain poorly understood.

## Results and Discussion

### Initial resting-state models of hH_V_1

We constructed three S4-down, resting-state models of the monomeric hH_V_1 VSD (residues M91–S224, the minimal functional sequence^19^). Based on the consistency of the R2 position and salt-bridge network with experimental data, two models were selected from those generated by the AF-Cluster approach^35,36^ and one from Robetta^42^ predictions (Methods, Figure 1b and Supplementary Figure S1). In these models, R2 is located just below the HG, and S4 is held in place through key salt bridges between the gating arginines and their counter-charges on S1 and S3. Specifically, R1 interacts with D112 (S1) and D185 (S3), while both R2 and R3 interact with D174 (S3); note, the R2–D174 salt bridge is absent in AF model 2. A comparison with existing models is given in Supplementary Table S1.

### WE-CpHMD captures the hHV1 activation pathway

Each of the three S4-down models was embedded in a solvated lipid bilayer composed of POPE and POPG lipids (3:1 ratio) and equilibrated in multiple stages for about 900 ns (Supplementary Table S2) using AMBER24 package.^43^ In the final 800 ns, the system was further relaxed using unrestrained all-atom CpHMD simulation^38,39^ at symmetric pH = 7. Next, CpHMD simulations with the WE protocol (WESTPA2.0^41^) were conducted under applied ΔV*_m_*=200 mV, pH_o_=7 and pH_i_=5 (Figure 1c), representing experimental conditions that favor channel activation. The progress coordinate was the S4 *z*-coordinate, defined as the center-of-mass *z* of R1–R2 (R205–R208) relative to that of S1–S3 backbone atoms. Details are in Methods. The WE-CpHMD simulations initiated from AF model 1 (WE-CpHMD run 1; aggregate sampling time of 19.8 *µ*s) revealed that S4 is displaced outward by one-click, accompanied by water permeation of the channel pore and proton titration (Figure 1d). We first discuss this set of WE-CpHMD simulations before turning to other simulations, which together totaled over 100 *µ*s (Supplementary Table S3).

#### One-click S4 displacement, salt-bridge reconfiguration, and water wires

Figure 2 and the accompanying Supplementary Figure S2 and S3 present an analysis of a representative continuous trajectory that resulted in one-click S4 displacement and formation of water wires. The latter is used to define channel activation and differentiate between activated and resting states. Comparing the time courses of the S4 *z*-coordinate and the key salt-bridge distances suggests two distinct conformational transitions prior to water permeation (Figure 2a). The first conformational transition occurs when the S4 *z*-coordinate increases by approximately 2 Å, accompanied by a reorientation of the R2 side chain (Supplementary Figure S2), which disrupts its salt bridge with D174 and moves it closer to D112 (Figure 2a). At the same time, the R2 sidechain approaches the level of the HG, as reflected by the R2 dihedral angle – defined as V177(C*β*)-F150(C*γ*)-V109(C*β*)-R208(C*ζ*)^19^ – decreasing to approximately 0*^◦^*(Supplementary Figure S2). A previous MD study demonstrated that this dihedral decreases from positive to negative as R2 moves above the HG.^19^ During these conformational changes, the other two key salt-bridges, R3–D174 and R1–D112, remain intact. In particular, the R1–D112 distance exhibits negligible fluctuation, indicating a stable resting state.

**Figure 2:**
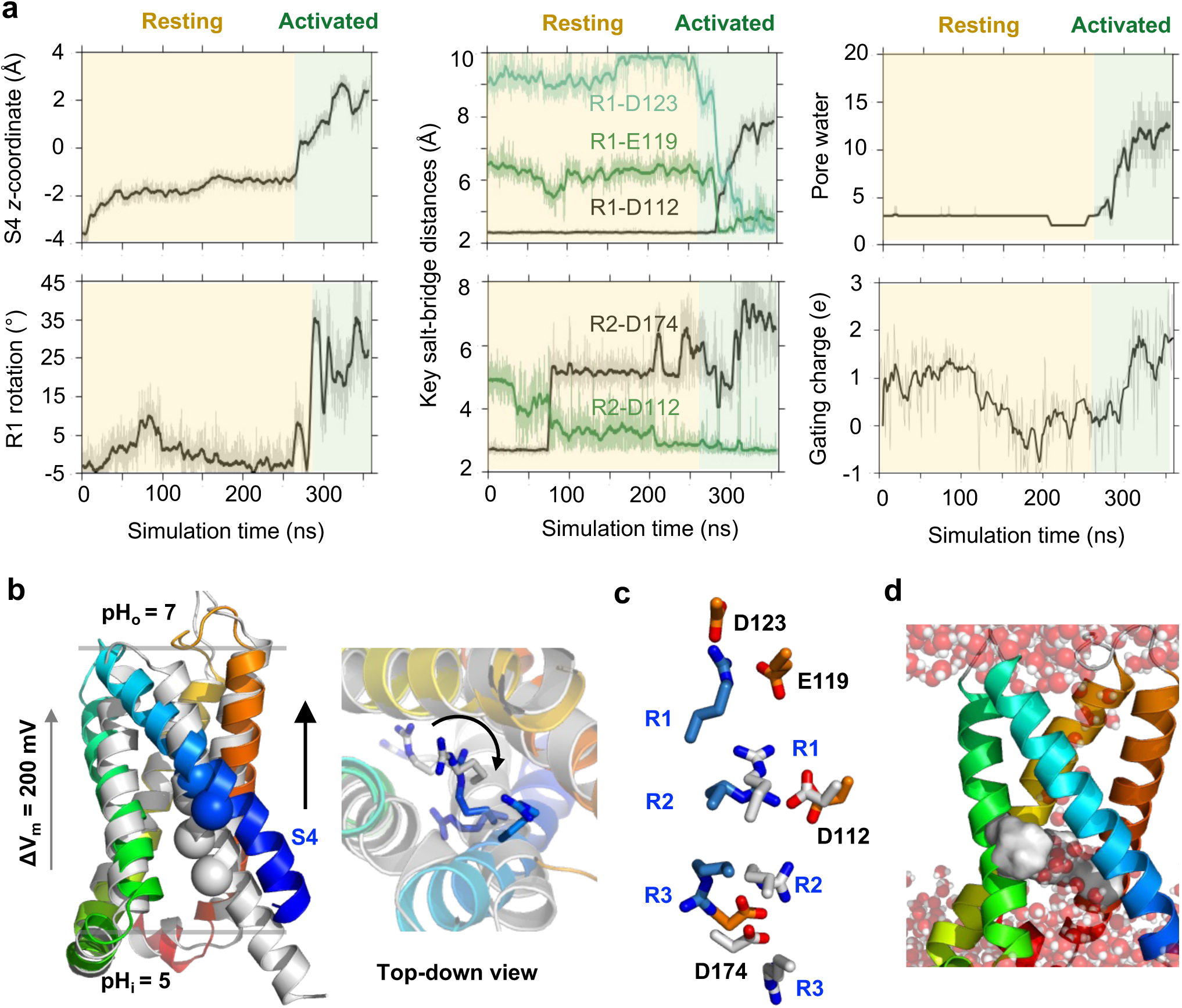
One-click conformational transition of hH_V_1 captured by the WE-CpHMD simulations under. Δ**V=200 mV and** Δ**pH=2.** A representative continuous trajectory initiated from AF Model 1 is analyzed. **a.** Time series of S4 *z*-coordinate, R1 rotation angle, distances of key electrostatic interactions involving R1/R2, the number of pore water, and the gating charge. S4 *z*-coordinate is defined as the center-of-mass displacement of residues between R1 and R2 relative to S1–S3. R1 rotation is defined as the angle between the C*α*–C*ζ* vector in the simulation frame and that in the initial frame. Pore waters refer to those within 3.4 Å of the D112 sidechain. Gating charge was calculated using the Q-route method. ^44^ Yellow and green shading denote the putative resting and activated states, respectively. **b.** Overlay of representative S4-up (rainbow) and S4-down (light gray) conformations. A top-down view is displayed, with R1/R2/R3 sidechains shown in blue (S4-up) and gray (S4-down). **c.** Zoomed-in view of key salt-bridge interactions in the S4-up (R1/R2/R3 in blue, acidic residues in orange) and S4-down (all residues in gray) conformations. **d.** A continuous water wire in the S4-up conformation. HG residues are displayed in surface rendering. For clarity, lipids are hidden. Additional analyses are given in Supplementary Figure S2 and S3.

This resting state persisted until a second conformational transition took place, whereby S4 moves up (outward) by over 4 Å and water molecules permeate the pore region (Figure 2a). Interestingly, the resulting water wires can be continuous or disrupted at R2 or R1 (Figure 2d and Supplementary Figure S3). Concomitantly, the most stable salt bridge between R1 and D112 is disrupted, leading to a rotation of the R1 sidechain, formation of the R1–E119 and R1–D123 salt bridges and breakage of the R1–D185 salt bridge (Figure 2a and Supplementary Figure S2). At the same time, R2 is moved above the HG (as indicated by the R2sc dihedral, Supplementary Figure S2), enabling it to establish a salt-bridge interaction with D112 (Figure 2c). This rearrangement of the salt bridges is in striking agreement with the one-click hypothesis.^13,26–28,33^ It is worth noting that R3 maintains electrostatic interaction and occasionally forms a salt bridge with D174 in the S4-up state (Supplementary Fig. S2).

#### S4 displacement remodels salt-bridge pairings and protonation states

In addition to the above analysis, we examined the complete WE-CpHMD dataset to investigate potential correlations between S4 displacement and reconfiguration of key salt bridges. The distributions of the S4 *z*-coordinate are plotted in the presence or absence of salt bridges characteristic of the resting or activated state of hH_V_1 (Figure 3a,b and Supplementary Figure S4a-c). Accordingly, the S4-up conformations (*z >* 0 as seen in Figure 2a, top left) are associated with the disruption of the R1–D112 and R2–D174 salt bridges (Figure 3a) and formation of the R1–E119, R1–D123, and R2–D112 salt bridges (Figure 3b and Supplementary Figure S4b). Moreover, the R1–D185 and R3–D174 salt bridges are largely incompatible with the S4-up state, whereas their disruption favors it (Supplementary Figure S4a, S4c). These data together suggest that the S4-up state is stabilized by the reconfigured salt-bridge pairings. The existence of the R1–E119 salt bridge in the activated H_V_1 supports Perozo’s hH_V_1 model based on the X-ray structure of activated Ci-VSD^28^ as well as the R2D model by Pomes, DeCoursey and coworkers.^13^

**Figure 3:**
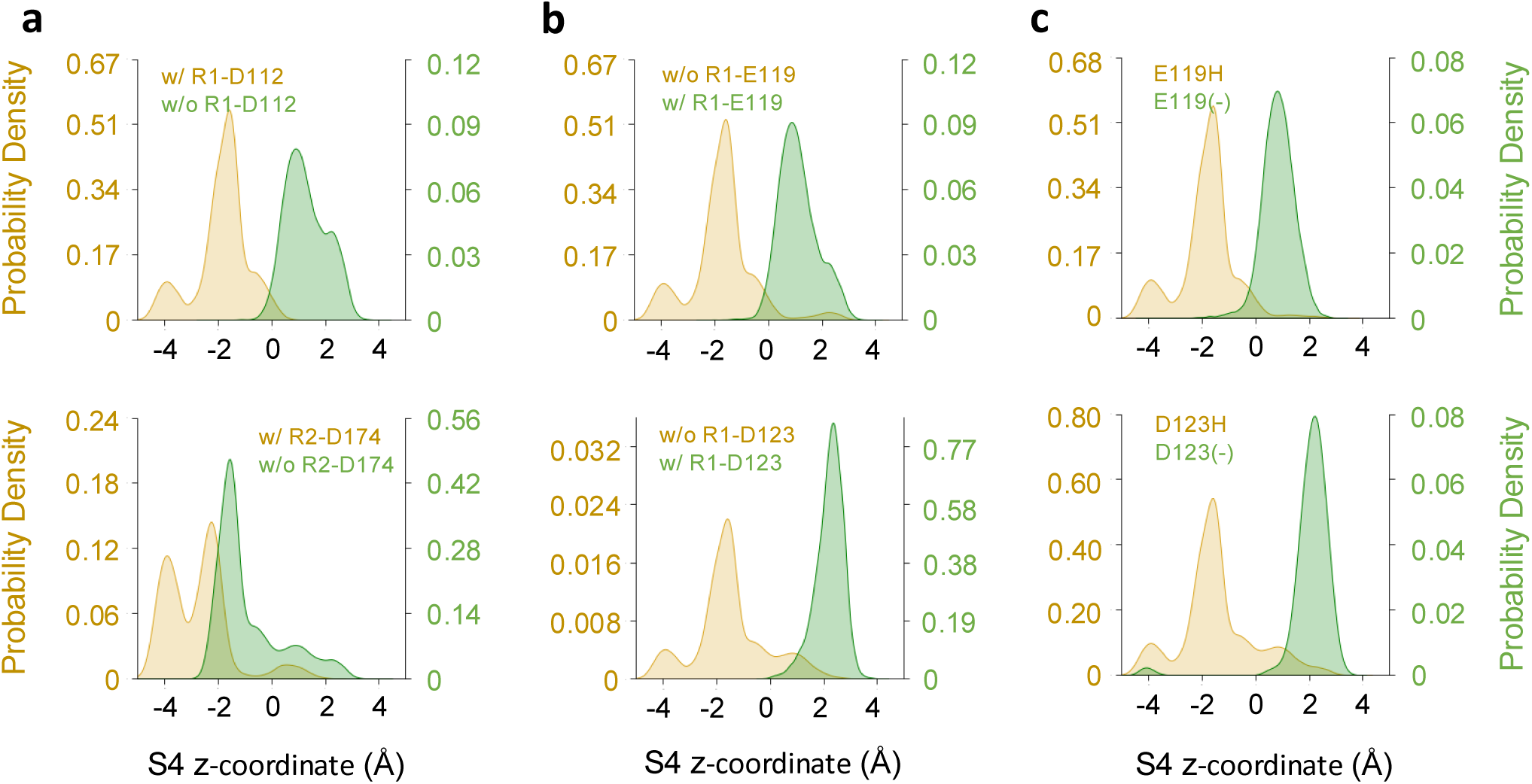
S4 displacement remodels salt-bridge pairings and protonation states in the WE-CpHMD simulations. Distributions of the S4 *z*-coordinate in the presence or absence of the specified salt-bridges or protonation state changes. The entire WE-CpHMD dataset initiated from AF model 1 is analyzed. The upward displacement of S4 is correlated with the disruption of R1–D112 and R2–D174 salt-bridges (**a**), the formation of R1–E119 and R1–D123 salt-bridges (**b**), and the deprotonation of E119 and D123 (**c**). A salt bridge is considered formed when the minimum carboxylate–guanidinium distance is less than 3.5 Å. Additional analysis is given in Supplementary Figure S4.

There are a total of 16 Asp/Glu/His residues in hH_V_1, nine exposed to the extracellular pH (7) and seven exposed to the intracellular pH (5). Only three residues – one facing the extracellular side (E192) and two facing the intracellular side (E164 and E171) – actively undergo titration, while several other residues occasionally switch their protonation states. (Supplementary Figure S2, bottom panels). Analysis of the complete WE-CpHMD dataset suggests that the deprotonation of E119 and D123 (protonated in the S4-down state) and the upward displacement of S4 are correlated (Figure 3c), This is attributed to the formation of R1–E119 and R1–D123 salt-bridge interactions, which in turn stabilizes the S4-up state.

### Comparison with X-ray structures of Ci-VSD, gating charge and sm-FRET measurements

To confirm the S4-down (putative resting state) and the S4-up (putative activated state) conformations of hH_V_1 obtained from the WE-CpHMD, two representative snapshots were selected to initiate extended, triplicate CpHMD runs (1.4 *µ*s each) under the same membrane potential and pH gradient. The S4-down state was further substantiated by prolonging the “equilibration” CpHMD simulation at pH 7, also performed as triplicate runs of 1.4 *µ*s each. Across the resting-state runs, no conformational rearrangement was observed. The S4 *z*-coordinate fluctuates around -4 Å, the HG remained closed, and gating arginines are involved in four salt bridges, R1–D112, R2–D174, and R3–D174/E171 (Figure 4a and Supplementary Figure S5). Across the activated-state runs, the S4 segment relaxed downward relative to the starting structure (S4 *z*-coordinate = 2.6 Å), with an average *z*-coordinate of 0.9 Å. Importantly, the characteristic salt bridges, R1–D123, R1–E119, and R2–D112 are intact (Figure 4a and Supplementary Figure S6).

**Figure 4:**
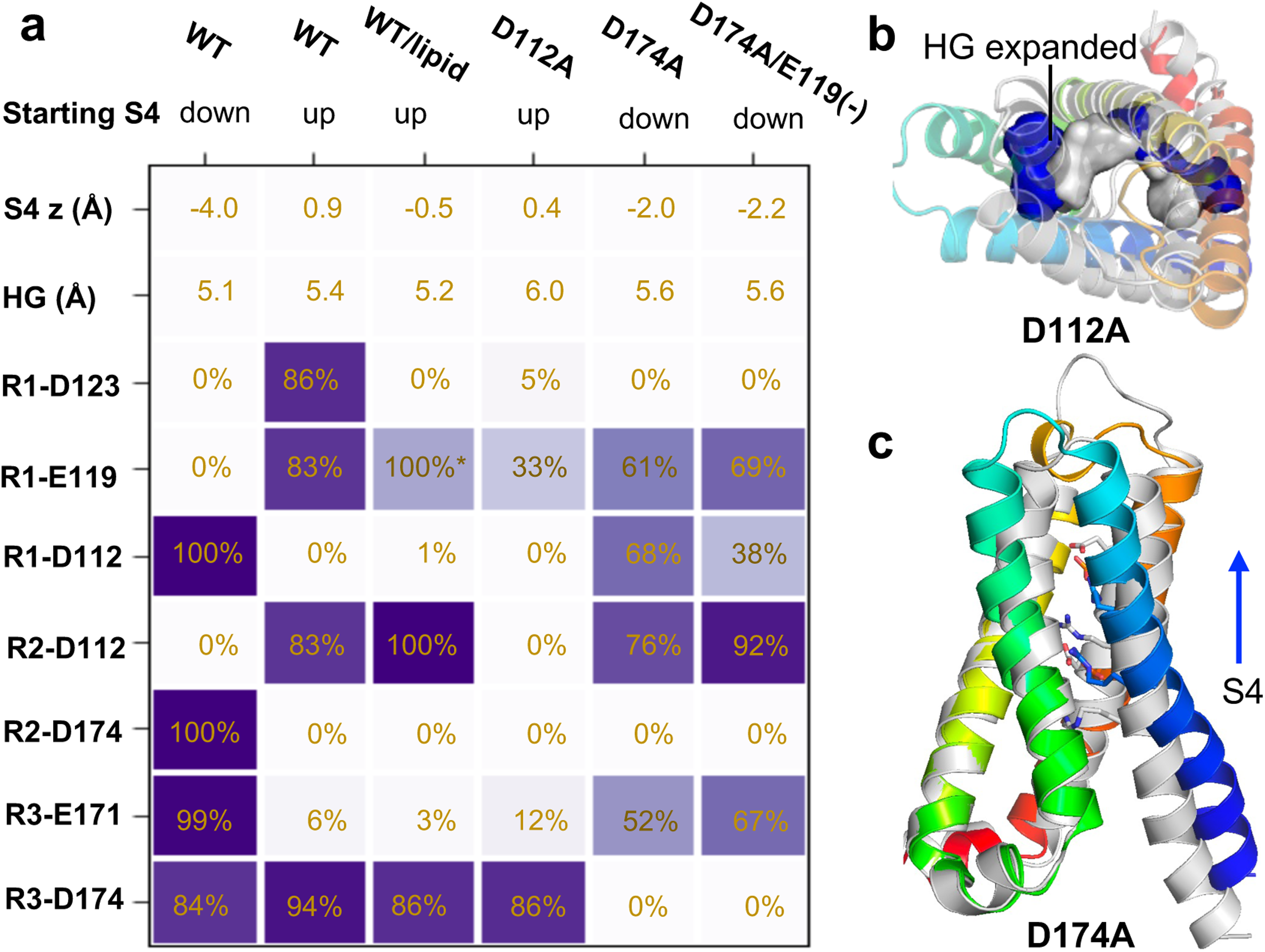
Characteristics of the S4-down and S4-up conformations of hH_V_1 and effects of D112A and D174A mutations. **a.** Average S4 *z*-coordinate, radius of gyration of the HG residues, and occupancies of key salt-bridge interactions in triplicate CpHMD simulations of WT, D112A, D174A, and D174A/E119(-), each initiated from either the S4-down or S4-up conformation. The last 500 ns of each trajectory were used for analysis. Data from the two WT runs in which lipid binding was observed are shown separately. Salt bridges are defined as in Figure 3. **b.** Top-down view of a representative snapshot from the D112A S4-up simulations (rainbow), superimposed on the WT S4-up structure (light gray), revealing an enlarged HG region, rendered as blue and gray surfaces for D112A and WT, respectively. **c.** A representative snapshot from the D174A S4-down simulations (rainbow) overlaid on the WT S4-down structure (light gray).

Following the extended simulations, we compared the putative resting and activated conformations of hH_V_1 to the corresponding X-ray structures of Ci-VSD.^33^ An overlay of the representative S4-down and S4-up conformations of hH_V_1, aligned using S1–S3, indicates that S4 shifts upward by one helical turn with Δ*z* = 4.7 Å, accompanied by a 2.7*^◦^* increase in the S3–S4 angle. These deviations are in remarkable agreement with those between the resting (PDB 4G80) and activated (PDB 4G7V) Ci-VSD structures,^33^ which show the S4 displacement of one helix turn, Δ*z* of ∼5.0 Å, and a 7.3*^◦^*-increase in the S3-S4 angle. Furthermore, the R1 and R2 sidechains in hH_V_1 are rotated clockwise by 35*^◦^* and 20*^◦^*, respectively, while the corresponding Ci-VSD structures show rotations of about 50*^◦^*. Note, the hH_V_1 simulations do not show R3 rotation, as the R3 remains engaged in a dynamic salt-bridge with D174. By contrast, R3 is rotated toward the pore in the X-ray structures of Ci-VSD.^33^

To further validate the magnitude of S4 displacement in hH_V_1, we compare the calculated and experimentally estimated gating charge, which represents the net electric charge displaced across the membrane upon channel activation. Gating charge can be determined by integrating the gating current; however, gating current measurement for H_V_1 is challenging because proton blockers have not been identified.^5,34^ Accordingly, the limiting slope of the conductance–voltage relationship is frequently used as an estimate.^34^ Based on this approach and assuming the monomer gating charge as half that of the dimer, the gating charge of monomeric hH_V_1 was estimated as 3.0^45^ (with an earlier work reporting a value of 0.9^2^), whereas a value of 2.05 was reported for mH_V_1cc (1.0 without the coiled-coil domain)^46^ and 1.05 for the N264R mutant of *Ciona*-H_V_1.^47^

We computed gating charge using the Q-route method,^44^ explicitly accounting for changes in protonation states (Methods). Tracking the S4 displacement, the calculated gating charge rises to approximately 2 *e*_0_ upon channel activation (Figure 2a, lower right panel). Since the coiled-coil domain, which was suggested to contribute a gating charge of 1.0 *e*_0_,^46^ is excluded from the hH_V_1 construct studied here, this value is in good agreement with the experimental estimate of 3.0 *e*_0_.^45^ Note that the transient drop in gating charge between 120 and 260 ns is attributable to the transient deprotonation of D130 (Supplementary Figure S2).

We further validate the above conformational transitions observed in simulations by comparison to a recent single-molecule fluorescence resonance energy transfer (sm-FRET) study.^48^ The measured FRET efficiencies for hH_V_1 dimer (on the N214R background) reconstituted into liposomes and labeled at the K125–S224 sites are 0.6 in the resting and 0.9 in the activated state.^48^ Using a recent computational protocol,^49^ we estimated FRET efficiencies of 0.78 and 0.86 for the resting and activated states of hH_V_1 monomer, respectively (Figure 1d). While the direction of the FRET change is consistent, the experimentally observed magnitude is substantially larger. This discrepancy may be attributed to the fact that the simulated hH_V_1 is a monomer lacking the coiled-coil domain. Furthermore, the observed FRET signal in the activated state may be overestimated due to contributions from signals between subunits.^48^

### Simulations of the D112A and D174A mutants

A vast number of mutations have been introduced in H_V_1 to probe the structure-function relationship.^34^ D112 is a selectivity-filter residue; neutral substitutions result in loss of proton selectivity or ion conductance.^30^ Consistently, in both the WE-CpHMD and extended CpHMD simulations, D112 forms the most stable salt bridge (Figure 2a and Figure 4a). The D112A substitutions in hH_V_1 shifted the activation threshold voltage by 59 mV, indicating a reduced propensity for channel activation.^12^ We conducted CpHMD simulations of D112A initiated from the S4-up conformation of WT hH_V_1, each lasting about 1.4 *mu*s. Although S4 remained in the up position with R2 above the HG, there was a significant backbone conformational change whereby the channel transitioned to an expanded state, as evident from a significant increase in the radius of gyration of the HG (Figure 4a,b and Supplementary Figure S6). This large conformational change is driven by the destabilization of the S1–S4 interactions mediated by the loss of the R2–D112 salt-bridge. Consequently, the R1–E119 and R1–D123 interactions were also abolished. These data suggest that, rather than returning the channel to the S4-down conformation, the D112A mutation causes the channel to expand. The loss of the salt bridge, the negative charge of D112, and the resulting expansion provides a plausible explanation for the observed chloride conduction by the D112A mutant.^30^

The D174A mutation was observed to negatively shift the threshold voltage of hH_V_1 by more than 100 mV,^12,50^ indicating an increased propensity for activation. To test this, we conducted simulations of D174A initiated from the S4-down conformation of WT hH_V_1. While S4 displacement is unlikely in such short (1.4 *mu*) simulations, one of the trajectories showed that D112 loses its interaction with R1 and instead forms the R2-D112 salt bridge characteristic of the activated channel. This likely results from the lack of the R2–D174 salt-bridge interaction, which frees R2 to engage with other acidic residues such as D112, thereby positioning its side chain above the HG (Figure 4a and Supplementary Figure S8).

### Effects of phospholipids on hH_V_1 from simulation and experiment

#### Lipid binding can both inhibit and promote S4 displacement

Curiously, the hH_V_1 channel did not open in WE-CpHMD run 3: the R1–D112 salt bridge remained intact and the HG stayed closed throughout the 15 *µ*s of simulations (Supplementary Figure S10). Analysis revealed that a POPE lipid inserted from the extracellular side and formed persistent h-bonds with protonated E119H, D123H, and Q194, as well as a salt bridge with H140 (Figure 5a). At the same time, R1 remained further away from E119 (mostly above 8 Å). Since the “productive” WE-CpHMD run 1 suggests that salt-bridge interactions between deprotonated E119/D123 and R1 are associated with S4-up displacement (Figure 3c) while experiments showed that zinc coordination with H140 inhibits H_V_1 acti-vation,^23,51^ we hypothesized that POPE coordination at this site exerts a similar inhibitory effect by blocking R1 from interacting with E119 and D123 and preventing S4 from moving up.

**Figure 5:**
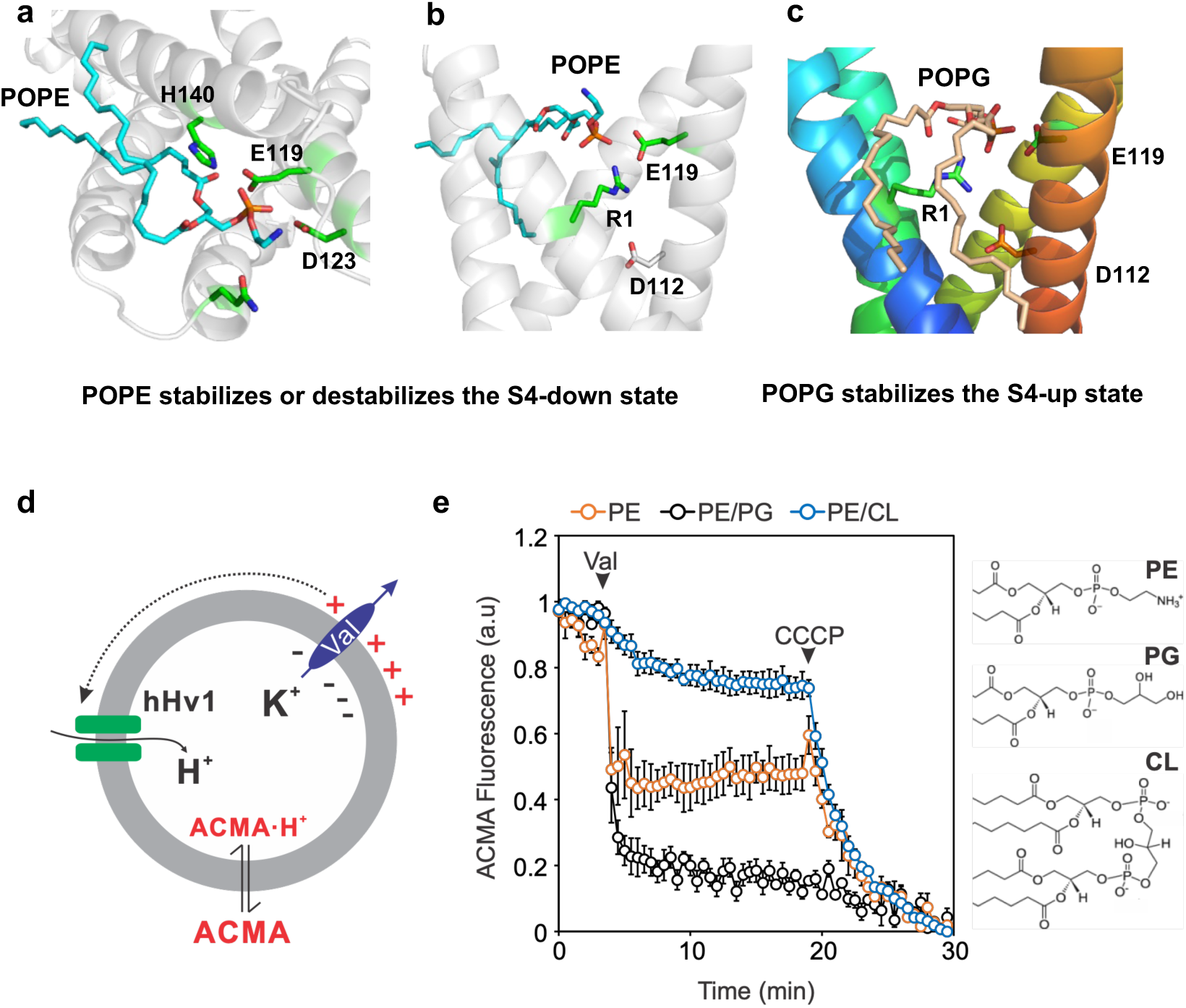
Lipid binding can inhibit or promote the activation of hH_V_1. **a.** A POPE lipid maintained stable h-bonds with E119H and D123H (S1), a salt bridge with H140 (S2), and a dynamic h-bond with Q194 (S3) throughout the WE-CpHMD Run 3 totaling ∼15 *µ*s, during which hH_V_1 remained locked in the S4-down conformation with the R1–D112 salt bridge intact (Supplementary Figure S9). **b.** POPE can also destabilize the S4-down state. In one D174A mutant run initiated from the S4-down conformation, a POPE lipid formed a long-lived (*>*1 *µ*s) h-bond with E119H and a salt bridge with R1, bringing R1 and E119H together, disrupting the R1–D112 salt bridge and promoting the S4-up displacement. **c.** POPG stabilizes the S4-up state. In two CpHMD runs initiated from the S4-up conformation, a POPG lipid bridged E119H and R1, analogous to POPE in **b**, stabilizing the S4-up state. **d.** Liposome flux assay monitoring proton uptake through hH_V_1 reconstituted into liposomes. A transliposomal *K*^+^ gradient in the presence of valinomycin establishes the membrane potential that drives proton uptake, reported as quenching of ACMA fluorescence. **e.** Liposome flux assay data for purified hH_V_1 reconstituted into liposomes, composed of 100% POPE (PE), POPE/POPG (3:1, PE/PG) or POPE/Cardiolipin (9:1, PE/CL). Assays were initiated by addition of valinomycin (Val, 0.45 *µM*). Carbonyl cyanide 3-chlorophenylhydrazone (CCCP, 1 *µM*) served as a positive control to confirm liposome integrity and the establishment of a positive outside liposome potential. Data are presented as mean±SEM (n = 3).

Interestingly, POPE simultaneously formed a hydrogen bond with E119H and a salt bridge with R1 in one of the D174A mutant runs initiated from the S4-down conformation. Around 550 ns, a POPE lipid inserted from the extracellular side and its phosphate group became coordinated with both E119H and R1. Concurrently, the critical salt bridge R1–D112 (hallmark of the closed state) broke, while S4 *z*-coordinate increased sharply (Supplementary Figure S8), indicating that channel opening was promoted. In contrast, another D174A mutant run showed that a POPG lipid became similarly coordinated with E119H and R1, although POPG-R1 interaction was significantly weaker, and R1–D112 salt bridge was maintained.

Bifurcated interactions between POPG and E119H/R1 were observed in two of three WT runs initiated from the S4-up conformation (Figure 5c and Supplementary Figure S6). Comparing to the WT run without POPG binding, the interaction between R2 and D112, another hallmark of the open state, is more stable (Figure 4a and Supplementary Figure S6). This suggests that POPG promotes or stabilizes the S4-up conformation by forming interactions with both E119H and R1, thereby bringing them in close proximity.

#### Effects of phospholipids on the function of hH_V_1

We reconstituted hH_V_1 into liposomes composed of POPE, POPE/POPG(3:1) or POPE/Cardiolipin (9:1) at a protein:lipid ratio of 1:200 and determined proton channel function using liposome flux assays.^48^ The membrane potential generated by a transliposomal *K*^+^ gradient via the ionophore valinomycin drives proton uptake into liposomes, leading to protonation and thus quenching of ACMA fluorescence (Figure 5d).

Although hH_V_1 channels in POPE liposomes are functional, their activity is markedly enhanced by the presence of 25% POPG (Figure 5e). Interestingly, cardiolipin, which, like POPG, carries a glycerol head group, has the opposite effect, strongly inhibiting proton uptake (Figure 5e). These functional data indicate that POPG enhances hH_V_1 channel function relative to POPE. This is consistent with the simulation data demonstrating that both lipids interact with counter-charges at the extracellular entrance of the hH_V_1 channel. While POPE binds the S4-down resting state – either stabilizing or destabilizing it – POPG binds and stabilizes the S4-up activated state.

### pH-modulated electrostatic breathing motions in the resting state

To further investigate electrostatic interactions and pH-dependent conformational dynamics of hH_V_1, we conducted equilibrium pH replica-exchange CpHMD^38,52^ simulations without transmembrane voltage or pH gradient. Simulations were initiated from the equilibrated Robetta model of S4-down hH_V_1 (Supplementary Figure S1) over pH 3.5–13.0, allowing all carboxylate and histidine sidechains to titrate, for a total of 7.2 *µ*s (Methods).

Across the experimentally relevant pH range of 4–8, the S4-down conformation persists, with the S4 *z*-coordinate spanning −5.5 to −3 Å and R2 remaining below the HG residues. Nevertheless, pH-dependent dynamics are observed in the salt-bridge interactions of the gating arginines, likely driven by sidechain rotamer adjustments (Supplementary Figure S11). For example, the R2–D174 salt bridge is significantly weakened at pH ≤ 6. These data indicate that in the absence of transmembrane potential or pH gradient, the channel is trapped within the S4-down conformational ensemble, exhibiting only localized, pH-modulated electrostatic breathing motions. Such pH-dependent dynamics facilitates the salt-bridge reconfiguration required for channel activation, e.g., weakening of the R2-D174 salt bridge upon intracellular acidification. Remarkably, of the salt bridges formed by the gating arginines, only R1–D112 remains intact across the pH range of 4–8 (Supplementary Figure S11).

In agreement with the WE-CpHMD and extended S4-down simulations, all histidines and a subset of carboxylates (E119, D123, D130, E153) have highly upshifted p*K* _a_’s and are fully protonated at physiological pH (Supplementary Figure S12). In contrast, D112 exhibits a “normal” solution p*K* _a_ below 4 (fully charged at physiological pH), which is consistent with the R1–D112 salt bridge being the most stable one in the WE-CpHMD simulations. This also aligns with its critical function in H_V_1, where a negatively charged carboxylate at this position is required for proton selectivity and channel gating^31^ (see also discussion of the D112A mutant). Notably, the outward-facing D185 and inward-facing D174 titrate in the physiological pH range, corroborating their flexible electrostatic interactions with the corresponding gating arginines observed in the WE-CpHMD simulations at pH*_o_*=7 and pH*_i_*=5. Protonation of D174 is responsible for the weakened interactions with both R2 and R3 as pH decreases below 7 (Supplementary Figure S12).

## Discussion

The WE-CpHMD simulations initiated from an experimentally consistent AF model reveal the voltage- and ΔpH-dependent activation pathway of hH_V_1. While supporting a one-click gating mechanism, the atomic details – including the putative resting and activated-state structural models – fill critical gaps in the structure-function understanding of hH_V_1 that are inaccessible to experiment, cMD, or direct AF modeling. Notably, none of the 812 AF models represents an open channel. Our simulations demonstrate that the transmembrane electric field sensed by the gating arginines triggers disruption of the R2–D174 salt-bridge interaction, allowing R2 to approach D112 and displace R1. This cascade breaks the linchpin R1–D112 interaction and establishes a new R2–D112 salt bridge, while the freed R1 shifts toward E119 and D123, forming salt bridges that stabilize S4 in the activated (up) position. The magnitude of S4 displacement (one helical turn), its tilting angle relative to S3, and salt-bridge reconfiguration are all consistent with the resting and activated Ci-VSD X-ray structures^33^ and the EPR-constrained hH_V_1 models.^28^ Accounting for the absence of the coiled-coil domain, the calculated gating charge of 2.0 is consistent with the estimated value of 3.0 for the monomeric hH_V_1 derived from the conductance-voltage relationship.^45^

Taken together, the extensive nonequilibrium (under applied membrane potential and pH gradient) and equilibrium CpHMD data suggest that the orchestrated conformational changes leading to hH_V_1 activation is facilitated by a delicately balanced electrostatic landscape and pH-modulated breathing motions in the resting state. While the selectivity-filter residue D112 remains deprotonated, locked in a persistent salt bridge with R1, other counter-charge residues such as D185 and D174 may protonate transiently, permitting breathing-like fluctuations of the salt-bridge network. Additionally, residues including E119 or D123 can deprotonate upon channel activation to stabilize the upshifted positions of the gating arginines. The use of a resting-state model based on the mH_V_1cc structure, together with the absence of dynamics-coupled proton titration inaccessible to cMD may account for the mechanistic discrepancy between the conclusions reached by the present work and a previous cMD study that proposed the two-click mechanism.^18^ Similarly, the CpHMD simulations refuted the earlier proposal that protonation of D112 allows the channel to open.^17,53^

The WE-CpHMD simulations demonstrate water permeation into the HG region and the formation of both continuous and disrupted water wires in the S4-up state. These data are compatible with the Grotthuss mechanism via a continuous water wire^12,20,54^ and a mechanism of proton hopping via both water and titratable side chains. The latter is conceptually similar to the recently proposed two-proton knock-on mechanism,^21^ although protonation of D112 itself is refuted by the present CpHMD data.

The present simulation work has several caveats. Kinetics of protonation events are not modeled by the CpHMD technique.^38^ The putative resting and activated-state models derived from the simulations may inherit certain inaccuracies from the initial AF model, e.g., the orientations of non-key residues may be less reliable. While a uniform constant electric field is an effective approximation of the membrane potential, it neglects spatial variations in the dielectric constant.^55^ The simulations applied different pH conditions to the inward- and outward-facing residues to mimic the pH difference across the membrane, an approach that is strictly valid only for the resting state. Finally, while the transmembrane pH difference is unaffected, the p*K* _a_ values of all residues are uniformly overestimated by up to two units due to the finite-size effect.^37,38^ These limitations are not expected to affect the mechanistic conclusions. Overall, this work demonstrates a computational framework broadly applicable to elucidating proton-dependent functional dynamics of proteins. This mechanistic principle – in which a delicately balanced electrostatic landscape and pH-modulated breathing motions in the resting state together prime the channel for voltage-gated opening – may underlie a broad range of proton-coupled biological processes.

## Methods

### Initial structural models of the resting-state hH_V_1

Structural predictions of the hH_V_1 channel were generated using AlphaFold (v2.3),^35^ AF-Sample,^56^ and AF-Cluster.^36^ AFSample and AF-Cluster are AlphaFold-based methods that each contain modifications of the pipeline to increase model diversity.^36,56^ Following the previous work, the hH_V_1 sequence (UniProt ID Q96D96) was truncated to the functional VSD region (residues 91–224), with the coiled-coil domain removed (termed minimal function sequence in Ref^19^). For AlphaFold, 20 random seeds, and no structural templates, were used to produce a total of 100 predictions.

AFSample was run with the “–model preset=monomer all” flag and v2.3 of AlphaFold, with all other settings at default. 600 structures were produced. For AF-Cluster, an MSA input file for the H_V_1 sequence was generated using ColabFold.^57^ The MSA was clustered using the default ClusterMSA.py script included in AF-Cluster, which produced 112 clusters. One model prediction for each cluster was generated using the AF-Cluster script run AF2.py, with no structure template.

We examined a total of 812 structural models generated by AlphaFold, AFSample, and AF-cluster for the presence of an intact HG and salt bridges characteristic of the S4-down, closed hH_V_1 channel (Figure 1a). The HG was considered intact if the pairwise distances between the sidechain center-of-mass of V109, F150, V177, and V178 were all below 4.5 Å. A salt bridge was considered formed if the minimum sidechain heavy-atom distance cutoff was below 3.5 Å. Only one model (AF model 1) satisfied all criteria – an intact HG and three salt bridges (R1-D112, R2-D174, and R3-D174), while a second model (AF model 2) met every criterion except R2–D174 (Supplementary Figure S1). These two models were energy minimized using the AMBER relax protocol^35^ before subject to MD simulations. It should be noted that none of the 812 models satisfies the criteria for an open hH_V_1 channel. Twelve models display an S4 *z*-coordinate greater than 2 Å; however, none exhibits the salt-bridge pattern characteristic of the activated S4-up state in either a one-click or two-click mechanism (Figure 1a).

We also generated an alternative closed-state model using the Robetta server^58^ with the full 273-amino acid hH_V_1 (HVCN1) FASTA sequence and no template. From the five predicted models, we manually selected the one consistent with the aforementioned structural features and extracted the aforementioned minimal functional sequence (residues 91–224)^19^ for pH replica-exchange CpHMD simulations.

### Molecular dynamics simulations

#### System preparation and equilibration

The simulation system was prepared with CHARMM-GUI.^59^ The capped protein (N-terminal acetylation and C-terminal amidation) was embedded in a 3:1 POPE/POPG bilayer (147 POPE, 49 POPG) using the PPM-generated orientation.^60^ A 15-Å-thick water layer was placed above and below the bilayer. 75 Na^+^ and 28 Cl^−^ ions were added to neutralize the system and reach an ionic strength of 0.15 M. The standard protonation states at pH 7, e.g., Asp(-), Glu(-), and His(0), were assumed.

The AMBER24 package^43^ was employed for all MD simulations. The protein was modeled with the CHARMM22 force field,^61^ including the CMAP correction;^62^ lipids with CHARMM36;^63^ and water with the CHARMM-modified TIP3P force field.^64^ Lennard-Jones and real-space electrostatic interactions were smoothly force-switched from 10 to 12 Å. Electrostatics were treated with the particle-mesh Ewald (PME) method^65^ using a reciprocal-space grid spacing of roughly 1 Å. Bonds involving hydrogen were constrained using the SHAKE algorithm.^66^ Following our previous protocol,^67^ the system was equilibrated in three stages. In the first two stages,^68^ restrained MD was performed with fixed protonation states (Asp(-)/Glu(-), His(0)) for 100.75 ns (Table S1). In the third stage, dummy hydrogen atoms were added to the Asp/Glu carboxylate and histidine imidazole groups, and the system was subjected to CpHMD equilibration at pH*_o_*=pH*_i_*=7 for 800.4 ns.

#### All-atom CpHMD simulations

Following equilibration, the system was subjected to weighted-ensemble (WE)-CpHMD simulations under a membrane potential of 200 mV and pH gradient of 2 units (pH*_o_*=7 and pH*_i_*=5) or pH replica-exchange (pHREX)-CpHMD simulations under symmetric pH and no membrane potential. Three sets of WE-CpHMD simulations were conducted, each initiated from the AF model 1 (run 1, 19.8 *µ*s), model 2 (run 2, 32.7 *µ*s), or the Robetta model of the closed hH_V_1 (run 3, 15.2 *µ*s). One set of pHREX-CpHMD equilibrium simulations was conducted starting from the Robetta model (7.2 *µ*s). Following the WE-CpHMD run 1, one S4-down and one S4-up snapshots were selected to initiate WT and mutant hH_V_1 simulations, each lasting 1.4 *µ*s (Table S3).

The membrane potential was represented^44^ by a uniform constant electric field of 0.0583 kcal mol*^−^*^1^ Å *^−^*^1^ e*^−^*^1^ across the simulation box (Figure 1d). To represent a transmembrane pH gradient, titratable residues were assigned with either intra- or extracellular pH based on their position relative to the HG (V109, F150, V177, and V178, Figure 1a). To impose a pH differential, titratable residues on the intracellular side were assigned a new titratable type in the CpHMD parameter file, in which the model p*K* _a_ values were increased by 2 pH units (while all other parameters were kept the same). Thus, inward-facing residues effectively experience a pH two units lower than outward-facing residues. The GPU-accelerated all-atom PME-CpHMD^37,38^ implementation was employed. All Asp/Glu/His residues were allowed to titrate, while the system net charge was compensated by the background plasma in the PME correction.^38^ All other CpHMD settings were kept at default values.^38^ The hydrogen mass repartitioning scheme^69,70^ was applied to allow a 4-fs timestep, with negligible impact on protonation states.^38^ The simulation was conducted in an NPT ensemble, maintaining the temperature at 310 K using a Langevin thermostat and the pressure at 1 atm using a Monte Carlo barostat.^43^

#### pH replica-exchange CpHMD

To characterize the protonation states of the closed hH_V_1 channel, equilibrium pH replica-exchange all-atom CpHMD simulations were conducted starting from the equilibrated Robetta structure. The simulations comprised 20 replicas, each assigned a distinct pH value (pH*_o_*=pH*_i_*) ranging from 3.5 to 11.0 in increments of 0.5. Conformational exchanges between neighboring pH replicas were attempted every 1000 MD steps according to the Metropolis acceptance criterion.^52^ Each replica was simulated in the NPT ensemble for 360 ns, with an aggregate sampling time of 7.2 *µ*s. All other setting were kept the same as in the WE-CpHMD simulations.

#### Weighted-ensemble (WE) CpHMD

To enhance sampling of rare events (S4-up displacement), we applied the weighted-ensemble (WE) protocol^71,72^ (implemented via the Python package WESTPA 2.0^41^) in the CpHMD simulations. The WE-CpHMD simulations were initiated from the AF model 1, model 2, or the Robetta model (Table S3). To capture the vertical displacement of S4 relative to the rest of the VSD, we defined the progress coordinate as the center-of-mass *z*-position of the R1–R2 side chains (R205, V206, A207, and R208) relative to that of S1–S3 residues (H99 to K125, Y134 to V160, and K169 to Q191), following alignment to the initial structure. The progress coordinate was first divided into bins with a width of 0.1 Å, and then manually refined to 0.05 Å to promote efficient displacement of S4. Each bin contained four walkers, and trajectories were propagated in segments of 100 ps resampling. During each WE iteration, walkers were split or merged as needed to maintain the target number of trajectories per bin while preserving statistical weights. The WE-CpHMD simulations were continued until water penetration of the pore was observed (along with S4 upshift of roughly one helical turn), for an aggregate sampling time of 19.8 *µ*s. No water penetration was observed in other two WE-CpHMD simulations starting from the AF model 2 or the Robetta model. To visualize the conformational changes occurring during activation, we generated a continuous transition trajectory by following and concatenating the sequence of walkers leading to the largest S4 displacement (Figure 2 and 3).

#### Extended CpHMD simulations of WT and mutant hH_V_1

From the WE-CpHMD simulation run 1, two representative structures corresponding to S4-down and S4-up states were extracted and used to initiate CpHMD simulations of the WT and three mutant (D112A, D174A, and D174A/E119(-)) hH_V_1 under the same membrane potential and pH gradient. Mutations were introduced using the PyMOL mutagenesis wizard.^73^ Mutant structures were minimized and equilibrated with position restraints using the same protocol as for setting up the CpHMD simulations. For each WT or mutant system, three independent trajectories (1.44 *µ*s each) were generated using different random velocity seeds (Table S3). HMR was applied in all simulations. The MD and CpHMD parameters were the same as those used in the WE-CpHMD simulations.

#### Gating charge calculations

To calculate gating charges, we used the Q-route method^44^ which expresses the excess charge as

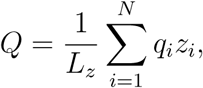

where *N* is the number of atoms, *q_i_* is the charge on atom *i*, *z_i_* is the *z*-coordinate of atom *i* in unwrapped coordinates, and *L_z_* is the length of the simulation box in *z* direction. The gating charge of between two conformations is given by *Q_g_* = *Q*_1_ − *Q*_2_. To account for protonation-state changes during channel activation, the net charge of a titratable residue was derived from the CpHMD *λ* value (protonated when *λ <* 0.2; deprotonated when *λ >* 0.8). Each residue’s net charge was then treated as a point charge located at its center-of-mass *z*-coordinate when evaluating the equation above.

#### Calculation of FRET efficiencies

The Python package FRETpredict^49^ was used to calculate FRET efficiencies. A recent smFRET study was conducted on the hH_V_1 dimer (N214R background) reconstituted into liposomes.^48^ To compare with this study, rotamers were placed on the smFRET labeling sites K125 and S224. To match the experimental donor and acceptor (Cy3 C5 and Cy5 C5 maleimide, respectively) as closely as possible, we used ‘Lumiprobe Cy3b C2R cutoff10’ as the donor rotamer library and ‘AlexaFluor 647 C2R cutoff10’ as the acceptor rotamer library, with a fixed Fö rster radius (*R*_0_) of 5.5 nm. Electrostatic calculations were enabled, the temperature was set to 298 K, and the dynamic regime was used.

#### Liposome fluorescence flux assay

hH_V_1 channels carrying an N-terminal 6xHis fusion tag were overexpressed in *E. coli* BL21 (DE3) host cells and purified by metal-affinity chromatography.^48^ The purified channels were reconstituted into liposomes containing different phospholipids at a protein-to-lipid ratio of 1:200 (w/w), with the reconstitution buffer containing 20 mM Hepes and 150 mM KCl, pH adjusted to 7.4 with 2.0 M NMDG. Liposome flux assays were conducted by diluting liposomes 10-fold with assay buffer containing 20 mM Hepes, 150 mM NMDG, and 2 *µ*M of the pH-sensitive fluorophore ACMA, pH 7.4. After recording baseline ACMA fluorescence for 4 min (Ex/Em = 390/460 nm), the assay was triggered by adding valinomycin (final concentration 0.45 *µ*M) to establish a positive outside-to-inside transmembrane potential, driving proton uptake into the liposomes. Quenching of ACMA fluorescence was monitored for 16 min, after which the proton ionophore CCCP (final concentration 1 *µ*M) was added as a positive control to confirm liposome integrity and the establishment of the outside-positive potential.

## Acknowledgment

Financial support by the National Institutes of Health (R35GM148261 to J.S., R01GM142816 to S.W., R35GM144083 to B.P., and R35GM151963 to T.E.D) is acknowledged.

## Data Availability

All simulation inputs, Python scripts for analysis, initial structural models, putative resting/activated models, and data used for all figures/tables can be found at https://github.com/JanaShenLab/Hv1. Trajectory files will be deposited in Zenodo.

## Supporting Information Available

Supporting Information contains 4 Supplementary Tables and 12 Supplementary Figures.

