## Supporting Information for "Activation pathway of the human voltage-gated proton channel"

@ *Joint first author*

### List of Tables


S1 Summary of structure models related to $hH_V1$ . . . . . . . . . . . . . . . . . . S-3

S2 Multi-stage equilibration protocol for $hH_V1$ . . . . . . . . . . . . . . . . . . . . S-4

S3 Summary of the MD simulations in this study . . . . . . . . . . . . . . . . . . S-5

S4 Result of WE simulation starting from Robetta model . . . . . . . . . . . . . S-6


### List of Figures


S1 Comparison of three initial S4-down, resting-state models of $hH_V1$ . . . . . S-8

S2 Supplementary analysis accompanying Figure 2 . . . . . . . . . . . . . . . . S-9

S3 Water wires in the S4-up state of $hH_V1$ . . . . . . . . . . . . . . . . . . . . . S-10

S4 Supplementary analysis accompanying Figure 3 . . . . . . . . . . . . . . . . S-11

S5 Simulations of $hH_V1$ starting from the S4-down conformation . . . . . . . . S-13

S6 Simulations of $hH_V1$ starting from the S4-up conformation . . . . . . . . . . S-14

S7 CpHMD simulations of D112A $hH_V1$ initiated from a S4-up conformation . . S-15

S8 CpHMD simulations of D174A $hH_V1$ initiated from a S4-down conformation S-16

S9 CpHMD simulations of the D174A/E119(-) $hH_V1$ initiated from a S4-down conformation . . . . . . . . . . . . . . . . . . . . . . . . . . . . . . . . . . . . S-17

S10 Distributions of key quantities in the WE-CpHMD run 3 . . . . . . . . . . . . S-18

S11 Key electrostatic interactions and S4 conformational properties from equilibrium CpHMD simulations . . . . . . . . . . . . . . . . . . . . . . . . . . . . S-19

S12 Titration of $hH_V1$ from equilibrium CpHMD simulations . . . . . . . . . . . . S-20

Table S1: Summary of published structure models related to hH$_V$1 in comparison to this work

| Model | Method | R1 partner | R2 partner | R2 position wrt. HG | R3 partner | |
|---|---|---|---|---|---|---|
| **S4-down (closed) state** | | | | | | **R3sc orient.** |
| mH$_V$1cc (3WKV)[S1] | X-ray | D112$^{(D108)}$ | None | Lower | D174$^{(D170)}$ | Lipids |
| Ci-VSD (4G80)[S2] | X-ray | D112$^{(D129)}$ | No | Same level | No | Lipids |
| Ci-VSD based hH$_V$1[S3] | EPR, Ci-VSD Ci-VSD (4G80) | D112, D185 | D174 | n/d | E153, E174 | Pore |
| Hypol hH$_V$1[S4] | mH$_V$1cc, cMD | D112, D185 | E153 | HG broken | None | Lipids |
| **This work** | WE-CpHMD | D112 | D174 | Lower | D174 | Pore |
| **S4-up (open) state** | | | | | | **S4 displ.** |
| Ci-VSD (4G7V)[S2] | X-ray | E119$^{(D136)}$ | D112$^{(D129)}$ | Higher | D174$^{(D186)}$ | One-click |
| | | | | R1sc/R2sc clockwise rotation | | |
| Ci-VSD based hH$_V$1[S3] | Ci-VSD (4G7V) | E119 | D112, D185 | Higher | No | One-click |
| Depol hH$_V$1[S4] | homology, cMD | D123 | E119, E192 | Higher | D112, D185 | Two-click |
| R2D hH$_V$1[S5] | homology, cMD | E119, D185 | D112 | n/d | D174, E153 | One-click |
| **This work** | WE-CpHMD | E119 | D112 | Higher | loosely D174 | One-click |
| | | | | R1sc/R2sc clockwise rotation | | |

Residue numbering displayed in the superscript corresponds to that in the PDB file. For the proposed models (CiVSD-based S4-down and S4-up models and the 'R2D' model), salt-bridge partners are adopted from the respective source publications. The 'R2D' model[S5] was obtained after 200 ns MD simulation starting from a homology model of hH$_V$1 using 3 templates (the VSD of K$_V$AP (PDB ID 1ORS), the paddle chimera of Kv1.2-Kv2.1 potassium channel (PDB ID 2R9R), and VSD of Na$_V$AB sodium channel (PDB ID 3RVY). The Ci-VSD based model for the S4-down hH$_V$1[S3] was obtained using homology modeling with the Ci-VSD structure (PDB ID 4G80) as a template and refined by PaDSAR approach after 10 ns MD simulation. The Ci-VSD based model for the S4-up hH$_V$1[S3] was proposed based on the one-click movement observed in the resting and activated Ci-VSD X-ray structures (PDB ID 4G7V, 4G7Y, and 4G80).[S2] The hypol and depol hH$_V$1 models were obtained from a MD trajectory[S4] initiated from a homology model based on the mH$_V$1cc X-ray structure (3WKV). The hypol model was obtained after $\sim$ 11 $\mu$s under -150 mV membrane potential. A subsequent 30-$\mu$s simulation under 150 mV membrane potential yielded the depol hH$_V$1 model. These membrane potentials were generated by applying a uniform electric field; however, the electric field strength was not specified in the paper.[S4] R1, R2, and R3 partners refer to the acidic residues forming salt bridges, defined using sidechain heavy-atom distance $\leq 3.5$ Å.

Table S2: Multi-stage equilibration procedure for $hH_V1$

| Stage | Ensemble | Timestep (fs) | Length (ns) | Restraints (kcal/mol/Å$^2$) Protein | Lipids |
|---|---|---|---|---|---|
| **Stage 1: Initial equilibration** | | | | | |
| 1.1 | NVT | 1 | 0.05 | 10 | 2.5 |
| 1.2 | NVT | 2 | 0.1 | 5 | 2.5 |
| 1.3 | NPT | 2 | 0.1 | 5 | 1 |
| 1.4 | NPT | 2 | 0.1 | 5 | 0.5 |
| 1.5 | NPT | 2 | 0.1 | 2.5 | 0.1 |
| 1.6 | NPT | 2 | 0.1 | 2.5 | 0 |
| **Stage 2: Membrane equilibration** | | | | | |
| 2.1 | NVT | 2 | 0.2 | 2.5 | 0 |
| 2.2 | NPT | 2 | 100 | 1.0 | 0 |
| **Stage 3: CpHMD equilibration at pH 7** | | | | backbone | |
| 3.1 | NPT | 2 | 0.2 | 1.0 | 0 |
| 3.2 | NPT | 2 | 0.2 | 1 | 0 |
| 3.3 | NPT | 2 | 800 | 0 | 0 |

Stage 1 follows the CHARMM-GUI protocol[S6] Unless specified otherwise, positional restraints on the protein were imposed on both backbone and side-chain heavy atoms. Restraints on lipids are applied to heavy atoms.

Table S3: Summary of the MD simulations in this study

| Simulation | Starting structure | Length (ns) | Voltage (mV) | $pH_o$ | $pH_i$ |
|---|---|---|---|---|---|
| **WE-CpHMD** | | | | | |
| WE1 | AF model 1 | 19,837 | 200 | 7 | 5 |
| WE2 | AF model 2 | 32,756 | 200 | 7 | 5 |
| WE3 | Robetta model | 15,165 | 200 | 7 | 5 |
| **Equilibrium pH replica-exchange CpHMD** | | | | $pH_o$=$pH_i$ | |
| REX | Robetta model | 360 x 20 | 0 | 3.5–13.0 | |
| **CpHMD** | Selected from the end of WE1 | | | | |
| WT S4-up | S4-up | 1,440 x 3 | 200 | 7 | 5 |
| WT S4-down | S4-down | 1,440 x 3 | 200 | 7 | 5 |
| D112A S4-up | S4-up | 1,440 x 3 | 200 | 7 | 5 |
| D174A S4-down | S4-down | 1,440 x 3 | 200 | 7 | 5 |
| D174A/E119(-) | S4-down | 1,440 x 3 | 200 | 7 | 5 |

Selection of starting structures is discussed in Methods. Both AF1, AF2, and Rosetta models contain R1-D112, R3-D174 and R2 is not higher than the hydrophobic gasket. The main difference is that AF model 1 and Rosetta model contains R2-D174 salt bridge, while AF model 2 does not. Results from AF1 model is used in the main text. Starting structures for the CpHMD simulations were selected from snapshots taken at the end of the WE1 simulations. The S4-down and S4-up conformation is selected based on the salt bridges and R2-HG relative positions listed in Table S1. WE2 simulations? Intracellular side lipid binding is observed in WE3 simulations.

Table S4: Result of WE simulation starting from Robetta model

| R1-E119 | R1-D185 | R1-D112 | R2-D112 | R2-D174 | R3-D174 | s4-$z$ | pore water |
|---|---|---|---|---|---|---|---|
| 2.7 | 7.8 | 10.1 | 2.7 | 6.9 | 2.7 | 4.5 | 4 |
| 3.5 | 10.9 | 12.5 | 2.8 | 7.3 | 2.7 | 5.1 | 4 |
| 3.0 | 12.0 | 11.9 | 2.8 | 7.3 | 2.7 | 4.6 | 6 |
| 2.6 | 11.8 | 10.8 | 2.6 | 6.9 | 2.7 | 5.6 | 4 |
| 2.9 | 8.8 | 10.4 | 2.8 | 6.2 | 2.7 | 5.2 | 4 |
| 2.7 | 12.2 | 13.2 | 2.7 | 7.3 | 2.7 | 4.4 | 7 |
| 2.6 | 6.3 | 8.2 | 2.6 | 6.9 | 2.7 | 4.4 | 7 |
| 2.7 | 5.7 | 7.9 | 2.7 | 6.1 | 2.7 | 3.2 | 6 |

Only 8 frames in WE meet the 'one-click' upshift of S4 helix. Pore water is defined as the number of water molecules within 3.4 Å of the D112 side chain.

### Supplementary Figures

#### 1. Initial S4-down $hH_V1$ models used in this work

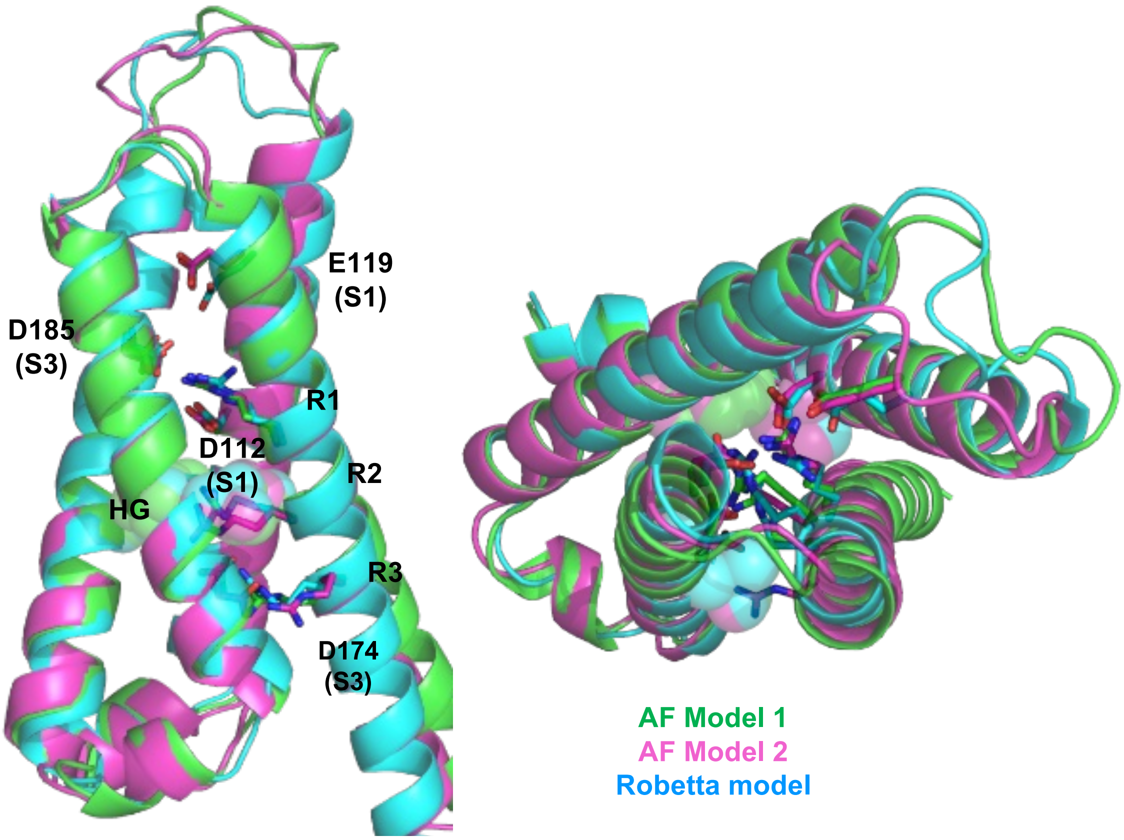


| | R1-E119 | R1-D185 | R1-D112 | R2-D112 | R2-D174 | R3-D194 |
|---|---|---|---|---|---|---|
| AF model 1 | 6.3 | 3.0 | 2.7 | 4.4 | 2.9 | 2.9 |
| AF model 2 | 6.3 | 2.9 | 2.8 | 5.3 | 5.2 | 2.7 |
| Robetta | 5.4 | 3.0 | 2.6 | 4.4 | 3.1 | 2.9 |

Figure S1: **Comparison of three initial S4-down, resting-state models of $hH_V1$.** Residues forming salt-bridge or electrostatic interactions with the gating charges (R1, R2, R3) are labeled, including D112 and E119 of S1, and D174 and D185 of S3. AF model 1 and model 2 were selected from the AFSampling and AF-Cluster protocol[S7,S8] (Methods). The Robetta model was selected from the five models downloaded from the server.[S9] In the WE-CpHMD simulations started from these three models, only the AF model 1 transitioned to the "activated" state. Salt bridges are identified in all three models and the sidechain distances are shown in Å. We define a S4 tilt angle with respect to S3 using vectors between the centers of mass of two sets of terminal residues on S4 (top: L198, G199, and L200; bottom: I218, S219, and V220) and S3 (top: D185, I186, and V187; bottom: I172, L173, and D174). After aligning the three models on the C$\alpha$ atoms excluding loops, the S4 tilt angle is 23.4°, 23.8°, and 24.3° in the AF1, AF2 (after energy minimization) and Robetta models, respectively. The C$\alpha$ root-mean-square deviations between AF model 1 and model 2, and between AF model 1 and Robetta model, are 0.89 and 1.41 Å, respectively.

#### 2. Additional analysis of the WE-CpHMD simulations

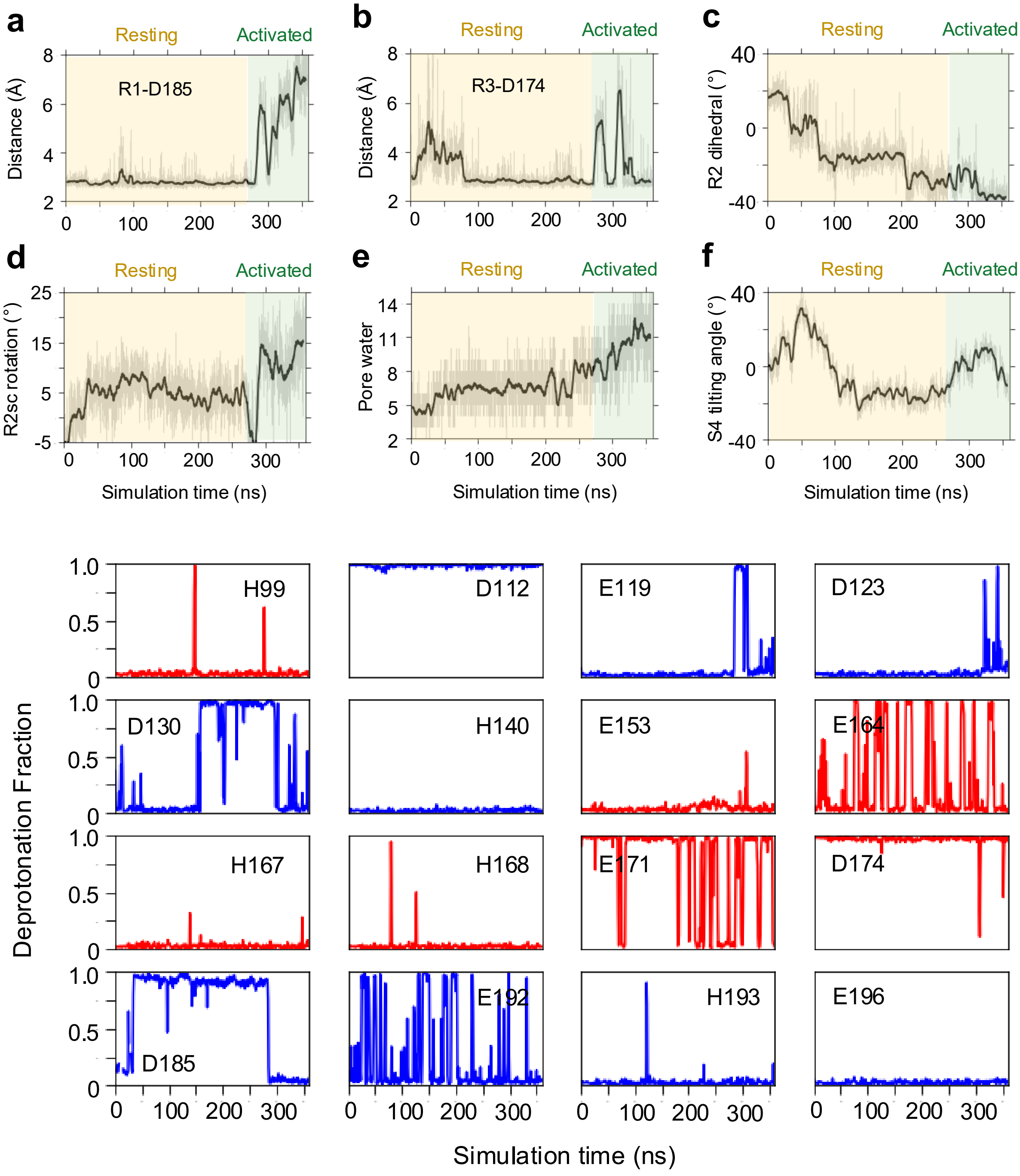


Figure S2: **Supplementary analysis accompanying Figure 2. Panels a-f**. Time series of minimum sidechain-to-sidechain distances (**a**, **b**), R2 dihedral angle representing its sidechain position relative to the HG, defined as V177(C$\beta$)–F150(C$\gamma$)–V109(C$\beta$)–R208(C$\zeta$) (**c**), R2 sidechain rotation angle, defined as the C$\alpha$–C$\zeta$ vector of the simulation frame relative to that in the initial frame (**d**), number of water with 3.4 Å to F150 (**e**), and S4 tilt angle as defined in Figures S1 (**f**). **Bottom four panels**. Time series of the deprotonation fractions of titratable residues. Deprotonation fraction is calculated over a time window of 1 ns. Deprotonated and protonated states are defined as $\lambda$ greater than 0.8 and less than 0.2, respectively. Residues exposed to the extra- and intracellular solution are colored blue and red, respectively.

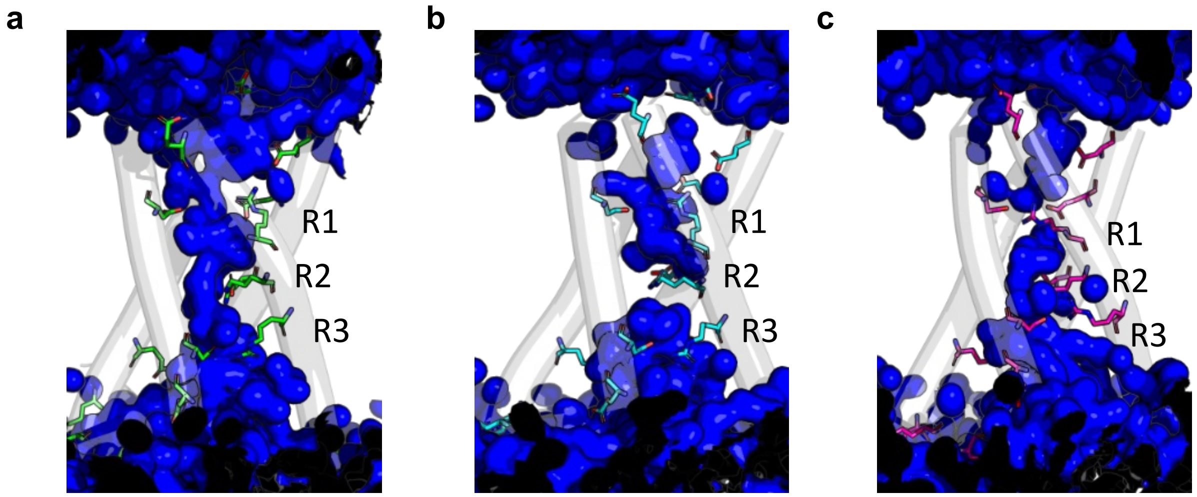


Figure S3: **Water wires in the S4-up state of hH$_V$1.** The water wire can be continuous (**a**) or interrupted at R2 (**b**) or R1 (**b**). Data taken from the representative continuous trajectory discussed in the main text and shown in Figure 2.

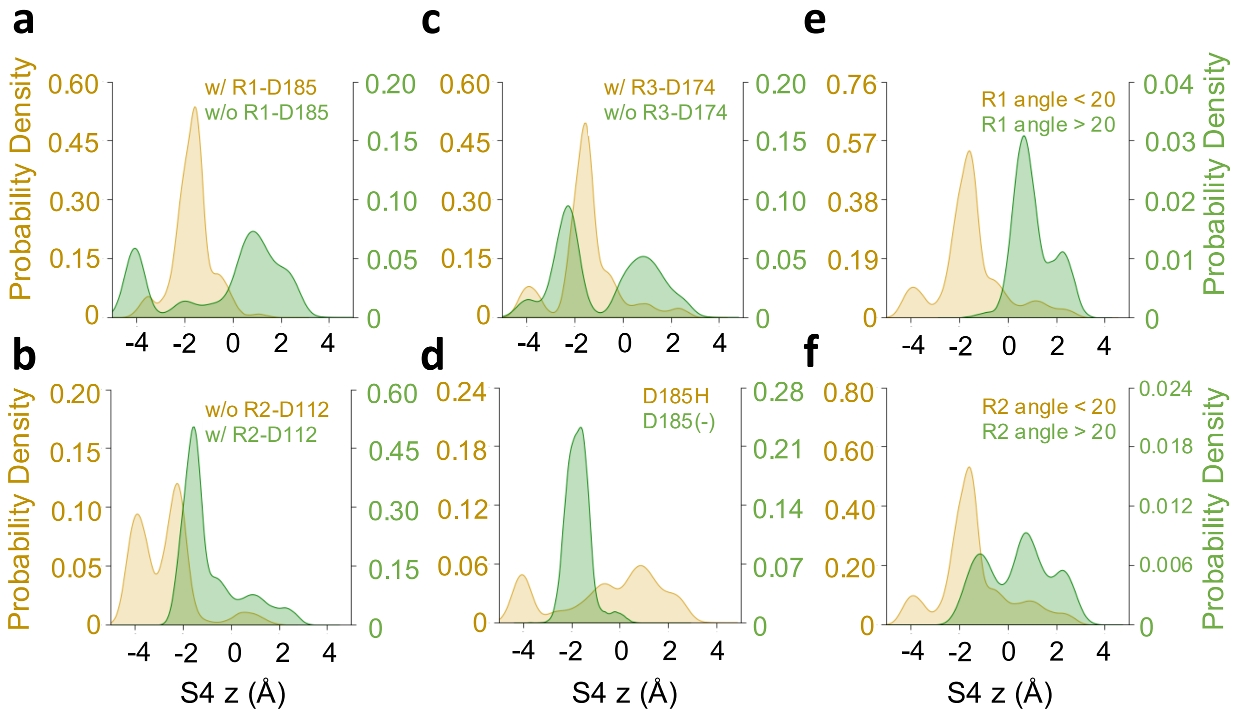


Figure S4: **Supplementary analysis accompanying Figure 3.** (**a.**) R1-D185, (**b.**) R2-D112, (**c.**) R3-D174, or in the protonated or deprotonated state of (**d.**) D185, or based on the rotation angle of (**e.**) R1, and (**f.**) R2. Salt-bridge is defined using a minimal sidechain-sidechain distance cutoff of 3.5 Å. The rotation angle is defined by the angle of the $C\alpha$–$C\zeta$ vector of Arg in the simulation frame relative to that in the initial frame.

#### 3. Analysis of the extended CpHMD simulations of WT and mutant $hH_V1$ initiated from a S4-down or S4-up conformation

Triplicate CpHMD simulations of $hH_V1$ initiated from both the S4-down and S4-up states were conducted under conditions of $V_m$ = 200 mV and $\Delta$pH = 2 ($pH_i$=5; $pH_o$=7). The distance for a salt-bridge interaction is defined as the minimal distance between side chain heavy atoms. The S4 $z$-coordinate is defined as the center-of-mass $z$ coordinate of R205–R208 residues relative to that of residues on S1, S2, and S3. The R2 dihedral angle, representing the vertical placement of its side chain relative to the hydrophobic gasket, is given by $C^{\beta}$(V177)– $C^{\gamma}$(F150)– $C^{\beta}$(V109) –$C^{\zeta}$(R208). An R2 value $\leq 0$ indicates that it lies above the hydrophobic gasket.[S10]

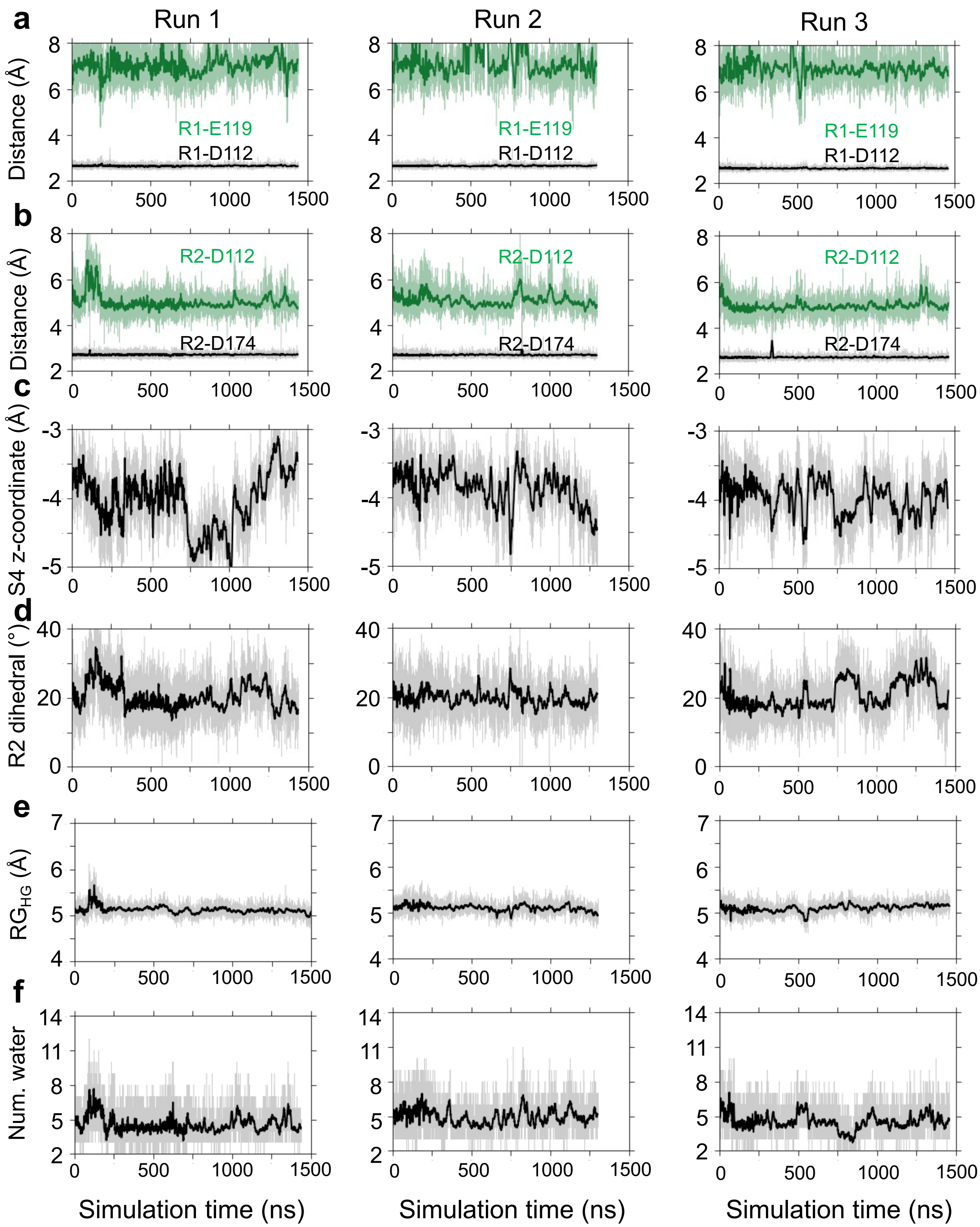


Figure S5: **Three independent CpHMD simulations of WT hH$_V$1 starting from the S4-down conformation.** $\Delta$V=200 mV and $\Delta$pH=2 was applied. Time evolution of the distances for **(a)** R1 and **(b)** R2 salt-bridge interactions, **c.** S4 $z$-coordinate, **d.** R2 dihedral angle, **e.** radius of gyration of HG, and **f.** the number of water within 3.4 Å to D112.

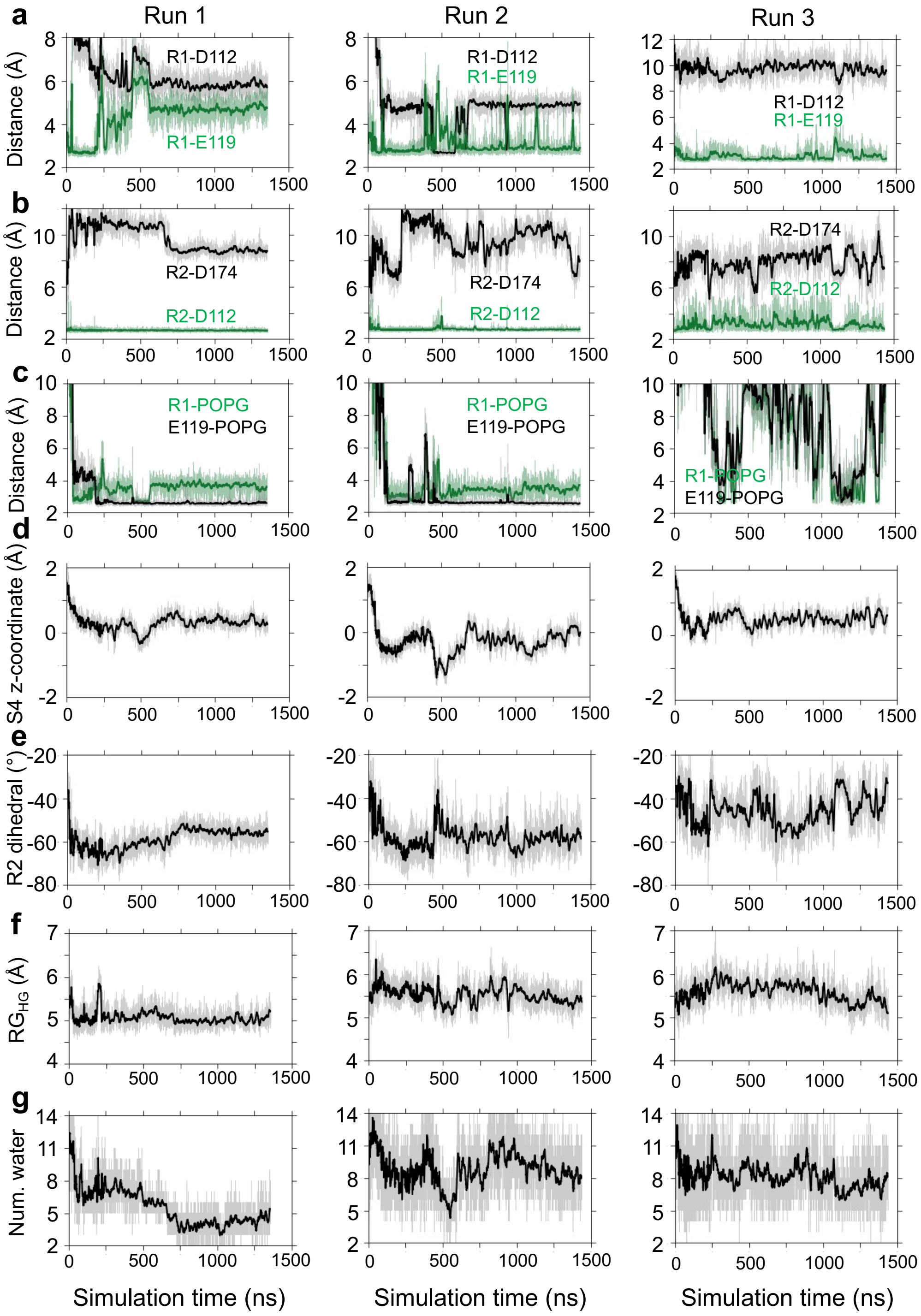


Figure S6: **Three independent CpHMD simulations of WT hH$_V$1 starting from the S4-up conformation.** $\Delta$V=200 mV and $\Delta$pH=2 was applied. Time evolution of the distances for **(a)** R1 and **(b)** R2 salt-bridge interactions, **d.** minimal distance between lipid to R1 and E119, **d.** S4 $z$-coordinate, **e.** R2 dihedral angle, **f.** radius of gyration of HG, and **g.** the number of water within 3.4 Å to D112.

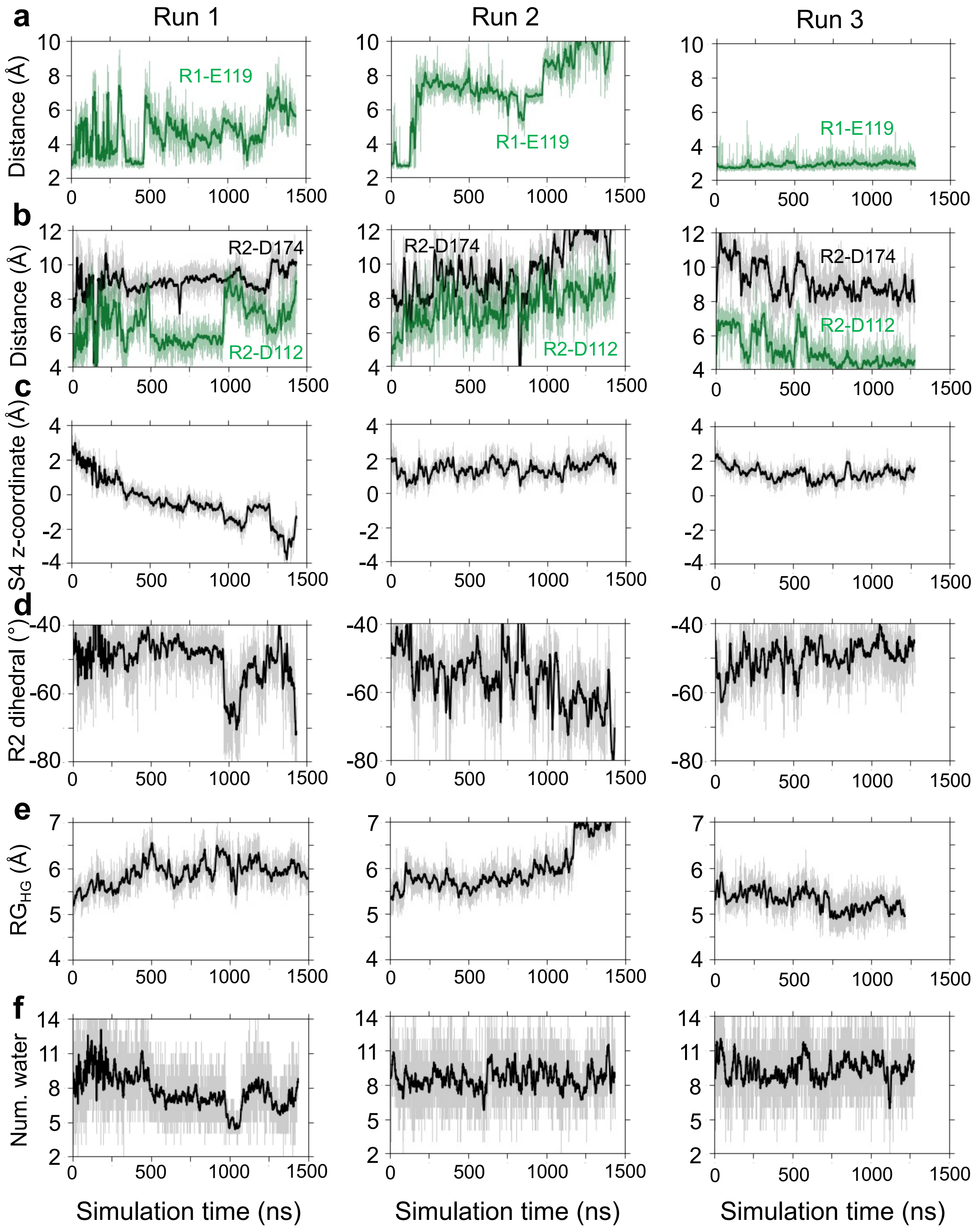


Figure S7: **Three independent simulations of D112A hH$_V$1 initiated from a S4-up conformation.** A membrane potential of 200 mV was applied in the presence of a transmembrane pH gradient (pH$_i$ = 5; pH$_o$ = 7). **(a)** R1 and **(b)** R2 salt-bridge interactions. **c.** Time evolution of the S4 $z$-coordinate, **d.** R2 dihedral angle, **e.** radius of gyration of HG, and **f.** the number of water within 3.4 Å to D112.

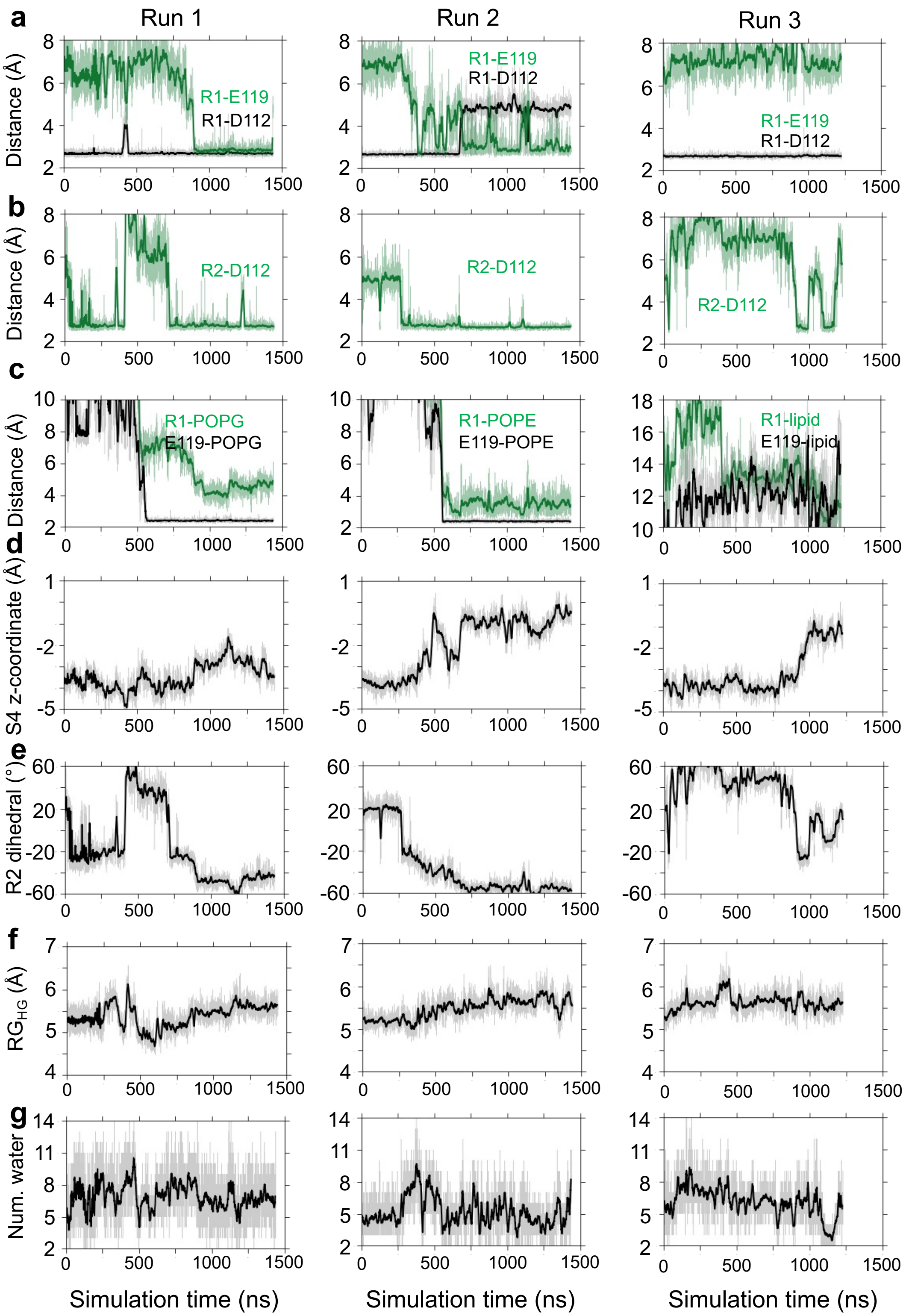


Figure S8: **Three independent CpHMD simulations of the D174A mutant $hH_V1$ initiated from a S4-down conformation.** A membrane potential of 200 mV was applied in the presence of a transmembrane pH gradient ($pH_i$ = 5; $pH_o$ = 7). Time evolution of the distances for **(a)** R1 and **(b)** R2 salt-bridge interactions, **d.** minimal distance between lipid to R1 and E119, **d.** S4 $z$-coordinate, **e.** R2 dihedral angle, **f.** radius of gyration of HG, and **g.** the number of water within 3.4 Å to D112.

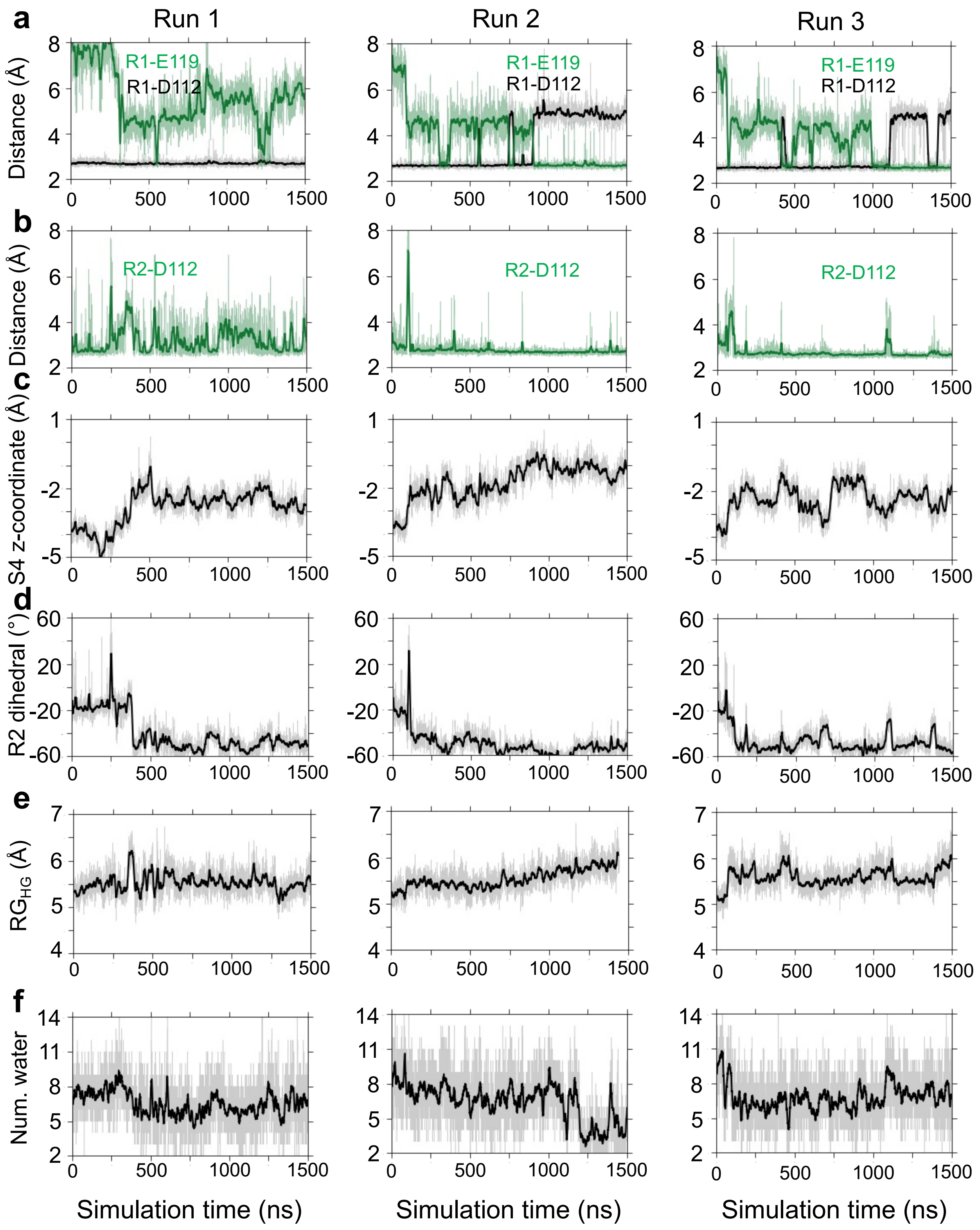


Figure S9: **Three independent simulations of D174A/E119(-) hH$_V$1 starting from the S4-down conformation.** A membrane potential of 200 mV was applied in the presence of a transmembrane pH gradient (pH$_i$ = 5; pH$_o$ = 7). **(a)** R1 and **(b)** R2 salt-bridge interactions. **c.** Time evolution of the S4 $z$-coordinate, **d.** R2 dihedral angle, **e.** radius of gyration of HG, and **f.** the number of water within 3.4 Å to D112.

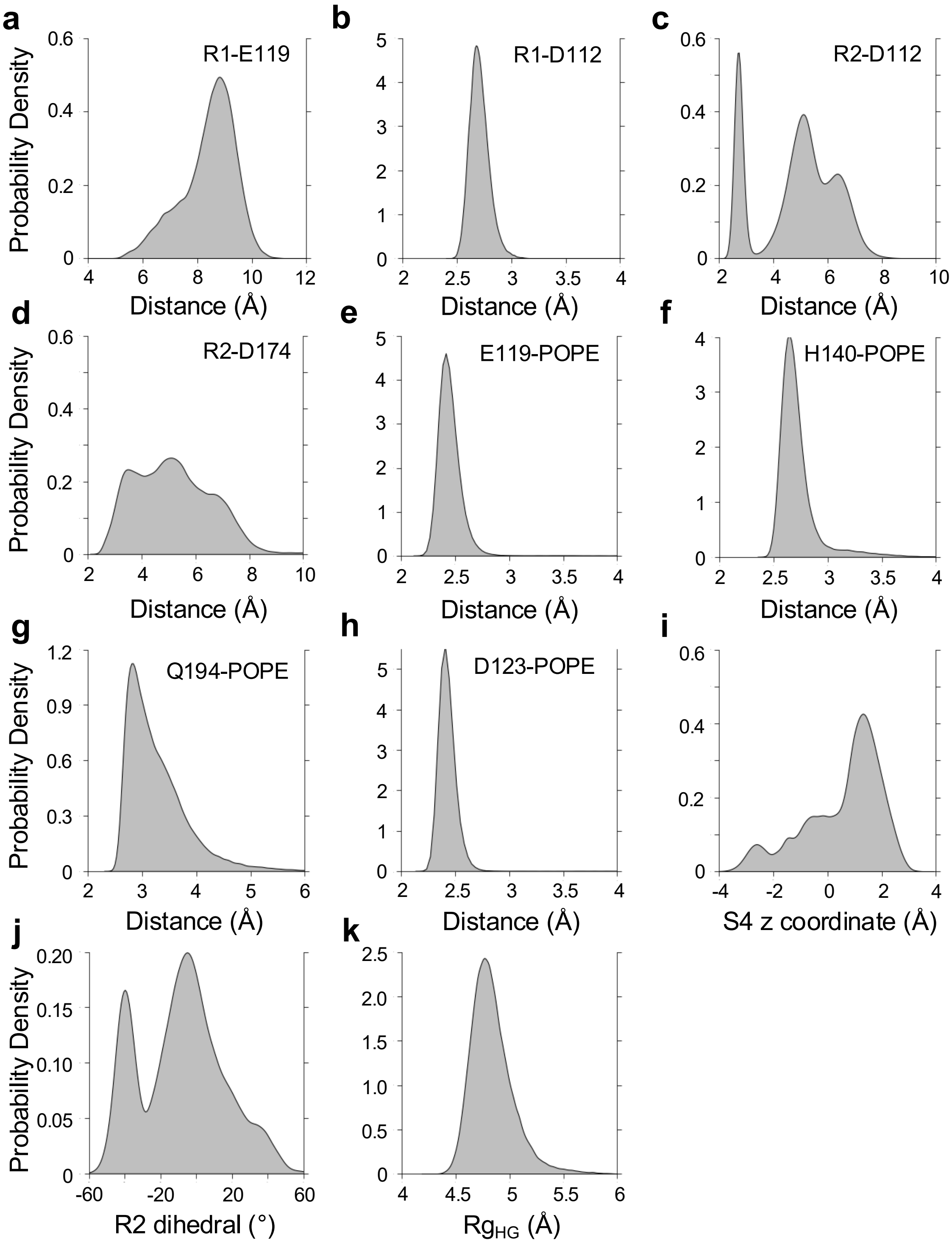


Figure S10: **Distributions of key quantities in the WE-CpHMD run 3 initiated from a Robetta S4-down hH$_V$1 model.** Distributions of the distances R1-E119 (**a**), R1-D112 (**b**), R2-D112 (**c**), R2-D174 (**d**), E119-POPE (**e**), H140-POPE (**f**), Q194-POPE (**g**), D123-POPE (**h**), and S4 $z$-coordinate (**i**), R2 dihedral angle (**j**), and gyration radius of the HG (**k**). Note: although S4 $z$-coordinate increased to nearly 2 Å and R2 dihedral angle became negative, R1–D112 stayed intact throughout the simulations, indicating that the channel remained closed.

#### 4. Equilibrium pH replica-exchange CpHMD simulations of $hH_V1$

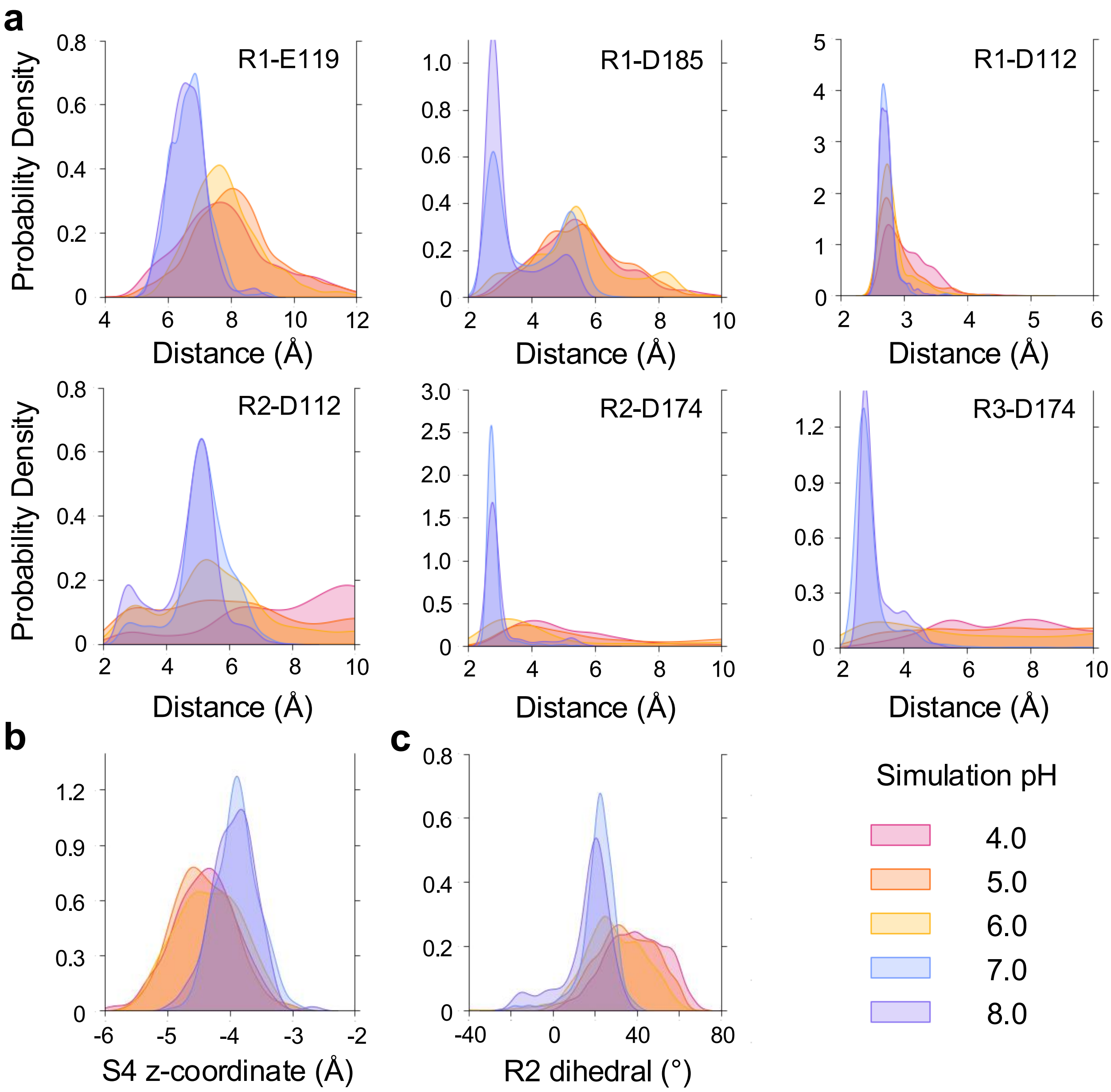


Figure S11: **Key electrostatic interactions and S4 conformational properties from CpHMD simulations at symmetric pH in the absence of membrane potential.** Distributions of the distances between the gating arginines and their counter charges (**a**), the S4 $z$-coordinate (**b**), and the R2 dihedral angle (**c**) at varying simulation pH. **Only one salt bridge, R1–D112, remains intact across the pH range 4–8.**

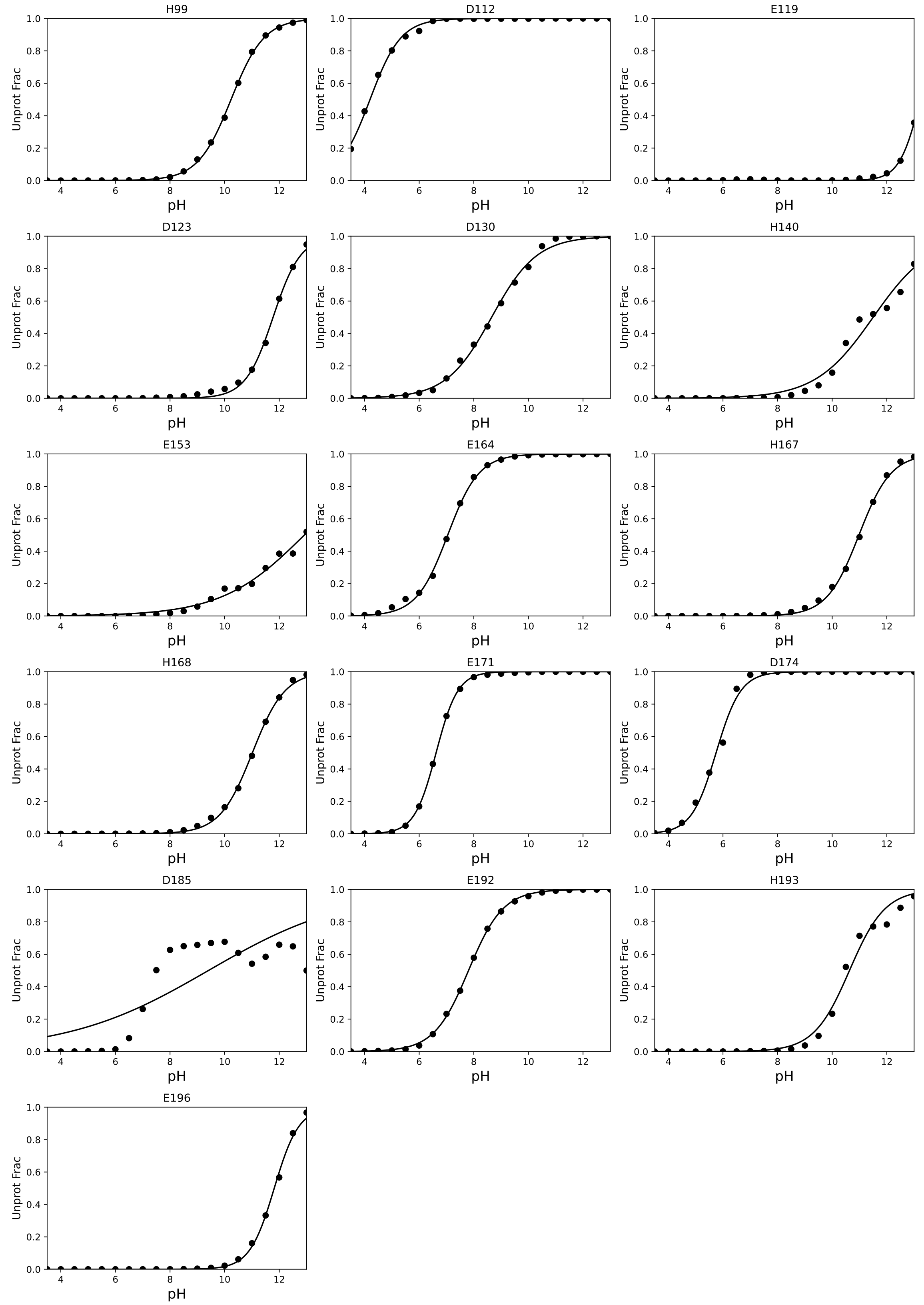


Figure S12: **Titration of $hH_V1$ from CpHMD simulations at symmetric pH in the absence of membrane potential.** Deprotonated fractions of Asp, Glu, and His sidechains in $hH_V1$ as a function of pH ($pH_o$ = $pH_i$, range 3.5–13 ) computed from the last 100 ns of the 360 ns (per pH replica) equilibrium CpHMD simulations.
